# ORP5/8 AND MIB/MICOS LINK ER-MITOCHONDRIA AND INTRAMITOCHONDRIAL CONTACTS FOR NON-VESICULAR TRANSPORT OF PHOSPHATIDYLSERINE

**DOI:** 10.1101/695577

**Authors:** Vera F. Monteiro-Cardoso, Leila Rochin, Amita Arora, Audrey Houcine, Eeva Jääskeläinen, Annukka M. Kivelä, Cécile Sauvanet, Romain Le Bars, Eyra Marien, Jonas Dehairs, Julie Neveu, Naima El Khallouki, Elena Santonico, Johannes V. Swinnen, David Tareste, Vesa M. Olkkonen, Francesca Giordano

## Abstract

Mitochondria are dynamic organelles essential for cell survival whose structural and functional integrity rely on selective and regulated transport of lipids from/to the endoplasmic reticulum (ER) and across the mitochondrial intermembrane space. As they are not connected by vesicular transport, the exchange of lipids between ER and mitochondria occurs at sites of close organelle apposition called membrane contact sites. However, the mechanisms and proteins involved in these processes are only beginning to emerge. Here, we show that the main physiological localization of the lipid transfer proteins ORP5 and ORP8 is at mitochondria-associated ER membranes (MAMs) subdomains, physically linked to the MIB/MICOS complexes that bridge the two mitochondrial membranes. We also show that ORP5/8 mediate non-vesicular transport of phosphatidylserine (PS) lipids from the ER to mitochondria by cooperating with the MIB/MICOS complexes. Overall our study reveals a novel physical and functional link between ER-mitochondria contacts involved in lipid transfer and intra-mitochondrial membranes contacts maintained by the MIB/MICOS complexes.

## INTRODUCTION

Vesicular trafficking is the major pathway for transport of proteins and lipids between membranes. However, an alternative route, which is vesicle-independent, occurs at regions of close inter-organelle membrane proximity (within less than 30 nm) also called membrane contact sites (Scorrano, De Matteis et al., 2019). This route is particularly important to preserve membrane composition, integrity and identity of intracellular organelles such as mitochondria that are largely excluded from the classical vesicle-mediated trafficking pathway. Like other organelles, mitochondria can be closely associated with the endoplasmic reticulum (ER), the major site of lipid synthesis and the major intracellular calcium (Ca^2+^) store. ER membrane subdomains closely apposed to mitochondria are called mitochondria-associated ER membranes (MAMs) and they facilitate the exchange of Ca^2+^ and lipids between the two organelles (Herrera-Cruz & Simmen, 2017, Tatsuta, Scharwey et al., 2014, Vance, 2014).

Mitochondria are involved in a plethora of cellular processes including energy production, lipid metabolism, Ca^2+^ homeostasis and apoptosis. To fulfill their numerous functions, mitochondria need to maintain a defined membrane composition by receiving essential lipids and lipid precursors from the ER through membrane contact sites (Acoba, Senoo et al., 2020, Giordano, 2018, Vance & Tasseva, 2013).

Increasing lines of evidence suggest that lipid transfer proteins (LTPs) play a major role in regulating the lipid composition of membranous organelles by facilitating non-vesicular lipid transport at membrane contact sites. In recent years, several tethering complexes with lipid transfer activity have been identified at membrane contact sites between the ER and other intracellular organelles as well as the plasma membrane (PM) in yeast and mammalian cells (Wong, Gatta et al., 2019). However, our knowledge of how lipids are exchanged at ER-mitochondria membrane contact sites remains rudimentary, and the LTPs that localize and function at these sites are just starting to be discovered. The best-studied lipid transfer/tethering complex at ER-mitochondria contact sites is the yeast ER-mitochondria encounter structure (ERMES) (Kornmann et al 2009, Lang et al 2015) that bridges the ER and the mitochondrial membranes and also facilitates the exchange of phospholipids (in particular phosphatidylcholine, PC) between them (ref). In metazoans, very little is known on how lipids are exchanged at ER-mitochondria membrane contact sites and about the proteins involved in this process. Some tethering proteins at mammalian ER-mitochondria contact sites have emerged in the recent years, such as VDAC-GRP75-IP3R and PTPIP51-VAPB complexes (Gatta and Levine 2017). A recent study has proposed that PTPIP51 could regulate cardiolipin levels by transferring its precursor phosphatidic acid (PA) to the mitochondria via its TPR domain (Yeo, Park et al., 2021). However, further work is required to clarify the lipid transfer ability of the PTPIP51 TPR domain (Francesca Giordano, 2021). Also, recently, mammalian LTPs with tethering function, such as VPS13A or Pdzd8 (the latter being proposed as a paralog of the ERMES subunit Mmm1(Wideman, Balacco et al., 2018)), were shown to localize at membrane contact sites, including those between ER and mitochondria, where they regulate membrane tethering and, in the case of Pdzd8, mitochondrial Ca^2+^ uptake (Hirabayashi, Kwon et al., 2017, Kumar, Leonzino et al., 2018). However, a direct involvement of these proteins in non-vesicular lipid transport between ER and mitochondrial membranes remains to be proved.

The Oxysterol binding protein (OSBP)-related proteins constitute a large family of LTPs conserved from yeast (Osh) to humans (ORP) and localized to different subcellular sites, shown in several cases to be membrane contact sites. A common feature of all ORPs is the presence of an OSBP-related lipid-binding/transfer (ORD) domain. Most ORP proteins contain a two phenylalanines (FF) in an acidic tract (FFAT)-motif that binds ER-localized VAP proteins and a pleckstrin homology (PH) domain that interacts with lipids or proteins in distinct non-ER organelle membranes. Two members of this family, ORP5 and ORP8, do not contain an FFAT motif but are directly anchored to the ER through a C-terminal transmembrane (TM) segment (Olkkonen, 2015).

ORP5 and ORP8 have been previously shown to localize at ER-PM contact sites where they transfer phosphatidylserine (PS) from the cortical ER to the PM, in counter-exchange with the phosphoinositides Phosphatidylinositol-4-phosphate (PI4P) and Phosphatidylinositol 4,5-bisphosphate (PIP_2_) (Chung et al. 2015; Ghai et al, 2017). We have recently shown that ORP5 and ORP8 are also present in the MAM and play a key role in maintaining mitochondrial integrity (Galmes, Houcine et al., 2016). We and others have also shown that ORP5/8 form a protein complex in the cell (Chung, Torta et al., 2015, Galmes et al., 2016). However, ORP5 and ORP8, when overexpressed, display a different distribution within MCS. In particular, overexpression of ORP5 greatly expands ER-PM contacts (Chung et al., 2015, Galmes et al., 2016), resulting in an accumulation of ORP5 at these sites, while overexpressed ORP8 is largely retained in the reticular ER. As all the studies on ORP5 and ORP8 so far have employed their individual overexpression, the endogenous sites where ORP5 and ORP8 interact and function as a complex are still unknown.

Interestingly, transport of PS is a key event occurring at ER-mitochondria contact sites. Newly synthesized PS, by the ER-localized PS-Synthase 1 (PSS1), is shuttled from the ER to the outer mitochondrial membrane (OMM) and from OMM to inner mitochondrial membrane (IMM) where it is rapidly converted to phosphatidylethanolamine (PE) by the PS-decarboxylase enzyme PISD (Tamura, Kawano et al., 2020, Vance, 2014, Vance & Tasseva, 2013). At the IMM, PE plays crucial roles in maintaining mitochondrial tubular morphology and therefore mitochondrial respiratory functions (Joshi, Thompson et al., 2012, Steenbergen, Nanowski et al., 2005). Regardless of extensive studies on PS transport between ER and mitochondria since its first discovery more than 20 years ago (Vance, 1990), the underlying mechanisms and proteins involved are still elusive.

Membrane contact sites exist also between the OMM and the IMM and are mediated by the Mitochondrial Intermembrane space Bridging (MIB) and Mitochondrial Contact sites and Cristae junction Organizing System (MICOS) complexes. The MICOS complex is a multi-subunit complex preferentially located at Cristae Junctions (CJ), tubular structures that connect the IMM to the cristae, and it is necessary for CJ formation, cristae morphology and mitochondria function (Harner, Korner et al., 2011, Huynen, Muhlmeister et al., 2016, Ott, Dorsch et al., 2015, Wollweber, von der Malsburg et al., 2017). The integral IMM protein Mic60 is the central component of the MICOS complex and carries a large domain exposed to the mitochondria intermembrane space (IMS) that interacts with the OMM Sorting and Assembly Machinery (SAM) to form the MIB complex (Friedman, Mourier et al., 2015, Guarani, McNeill et al., 2015). The SAM complex is constituted of SAM50 (a pore-forming β-barrel protein), metaxin1 and 2, and is involved in the membrane insertion and assembly of mitochondrial β-barrel proteins (Hohr, Lindau et al., 2018, Kozjak, Wiedemann et al., 2003, Kozjak-Pavlovic, Ross et al., 2007). However, whether and how OMM-IMM contact sites are linked to ER-mitochondria contacts in mammalian cells is still largely unknown.

Here we uncover for the first time the endogenous localization of ORP5 and ORP8, revealing that the major site of their interaction in physiological conditions are the MAMs. We also show that the ER subdomains where ORP5 and ORP8 reside are physically connected to the intra-mitochondrial membrane contacts bridged by the MIB/MICOS complexes at cristae junctions. We then show that ORP5/8 cooperate with SAM50 and Mic60, key components of the MIB/MICOS complex, to mediate PS transport from the ER to the mitochondrial membranes at ER-mitochondria contact sites in mammalian cells, and consequently the synthesis of mitochondrial PE.

Our findings reveal a novel tripartite association between the ER and the two mitochondrial membranes that links lipid transfer across these membranes, cristae biogenesis and consequently mitochondria function.

## RESULTS

### ER-mitochondria contact sites are the main localization of the ORP5-ORP8 complex

The localization of ORP5 and ORP8 at endogenous level is still unknown. We thus investigated their endogenous localization by immunofluorescence (IF) using antibodies against ORP5 and ORP8 proteins. First, we validated the specificity of these antibodies in cells overexpressing ORP5 or ORP8 proteins fused with a similar fluorescent tag (EGFP-ORP5 or EGFP-ORP8) and found that ORP5 and ORP8 signals detected using these antibodies co-localized with the overexpressed proteins (Fig. S1a-b). Then, we analyzed ORP5 and ORP8 endogenous localization in control HeLa cells and in cells where ORP5 and ORP8 were downregulated by RNAi and whose mitochondria were labeled by MitoTracker. We found a strong decrease in ORP5 and ORP8 IF labeling upon their knockdown (KD) (Fig. 1a, 1c, S1f), whose efficiency was confirmed by western blotting (WB) (Fig. 1b, S1c-f), validating the specificity of the used antibodies. ORP5 and ORP8-positive compartments in control conditions overlapped with the ER protein RFP- Sec22b, confirming endogenous ORP5 and ORP8 localization to the ER and further validating the specificity of these antibodies (Fig. S2a). Interestingly, the majority of endogenous ORP5 and ORP8 co-localized to subcellular compartments in close proximity to mitochondria (Fig. 1a, 1d, 1e-f).

**Figure 1.**
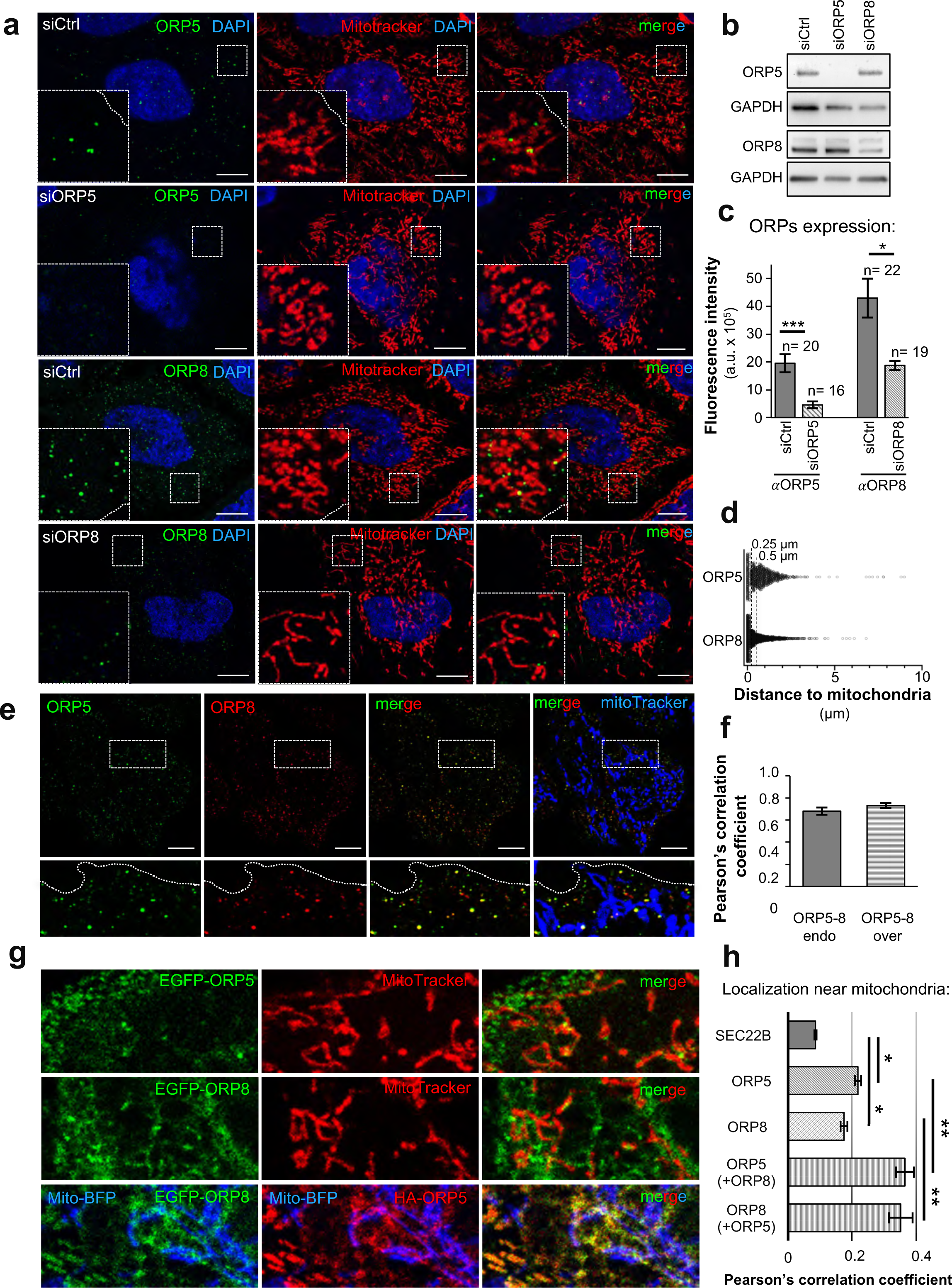
Endogenous and co-overexpressed ORP5 and ORP8 co-localize at ER-mitochondria contact sites. (a) Confocal images of Ctrl, ORP5 and ORP8 knockdown HeLa cells were immunostained using anti-ORP5 or ORP8 antibodies (green), and treated with MitoTracker to label mitochondria (red) and DAPI to stain the nuclei (blue). Images are presented as individual layers. Insets show magnifications of the boxed regions. Note the close association of endogenous ORP5 and ORP8 to mitochondria. Scale bar, 10 μm. (b) WB analysis showing ORP5, ORP8 and GAPDH levels in protein lysates from Ctrl, ORP5 and ORP8 knockdown HeLa cells. (c) Quantification of ORP5 and ORP8 fluorescent intensity in Ctrl, ORP5 and ORP8 knockdown cells. Mean of fluorescent intensities in arbitrary units (a.u. x10^5^). Error bars denote ± standard error of the mean (SEM). Number of cells given above bars. Statistical analysis: P values were determined by unpaired student’s t-test, *P<0.05, ***P<0.001. (d) Distribution of ORP5 and ORP8 IF staining (spots) in relation to their distance (in µm) to mitochondria indicating that part of the endogenous ORP5 and ORP8 in the cell is detected in close proximity to mitochondria (<0.5 µm). (e) Confocal images of a HeLa cell immunostained using anti-ORP5 (green) or ORP8 (red) antibodies and MitoTracker (blue). Images are presented as individual layers. Insets show magnifications of the boxed regions. Scale bar, 10 μm. (f) Quantification of the co-localization (Pearson’s factor) of ORP5-ORP8 in endogenous (ORP5-8 end) and co-overexpression (ORP5-8 over) conditions. Bars indicate mean values ± SEM. Number of cell analyzed: ORP5-8 end (n=15), ORP5-8 over (n=14). (g) Confocal micrograph of a region of HeLa cell (zoomed from Fig EV1a) transfected with EGFP-ORP5 (green), EGFP-ORP8 (green) or EGFP- ORP8 (green) + HA-ORP5 (anti-HA, red), and with Mito-BFP (blue). (h) Quantifications of the association to mitochondria (Pearson’s factor) of the indicated EGFP-tagged constructs. Bars indicate mean values ± SEM of three independent experiments with 10 cells for sample analyzed in each experiment (n=30). Statistical analysis: unpaired Student’s t-test comparing EGFP-ORP5 (ORP5) or EGFP-ORP8 (ORP8) to EGFP- Sec22b (SEC22b) and HA-ORP5 (+EGFP-ORP8) or EGFP-ORP8 (+HA-ORP5) to EGFP-ORP5 or EGFP-ORP8, respectively. *P<0.05, **P<0.01.

We then sought to analyze ORP5 and ORP8 localization when co-overexpressed (at similar levels) by co-transfecting HA-ORP5 and EGFP-ORP8 in HeLa cells and comparing their localization with the individually expressed EGFP-ORP5 and EGFP-ORP8 by confocal microscopy. When expressed alone, EGFP-ORP5 localizes to ER in contact with mitochondria, but also strongly increases ER-PM contact sites where it relocates, while it localizes very little in the reticular ER (Fig. 1g, S2b and (Galmes et al., 2016)). Instead, EGFP-ORP8, when expressed alone, localizes mostly to ER-mitochondria contacts and to reticular ER, with only a minor pool at cortical ER, as it does not increase ER-PM contact sites (Fig. 1g, S2b). Remarkably, even if the individually overexpressed ORP5 and ORP8 were differently distributed among MCS, their localization at ER-mitochondria contacts was higher as compared to a general ER protein, such as Sec22b (Fig. 1h). Moreover, and interestingly, when expressed together, ORP5 and ORP8 equally redistributed and co-localized to cortical ER, reticular ER and ER-mitochondria contacts (Fig. 1g, S2b). In particular, their localization to ER-mitochondria contacts was higher as compared to when expressed individually (Fig. 1h). Also, the co-localization of co-overexpressed ORP5 and ORP8 was comparable to the co-localization of the endogenous proteins, as revealed by the high Pearson’s correlation coefficient that was in both cases close to 1 (Fig. 1f). These data indicate that co-expression of ORP5 and ORP8 mimics the physiological localization of these proteins as a complex, when the expression levels of one of the two proteins are not highly enriched as compared to the other.

To further quantify ORP5 and ORP8 co-localization and interaction at ER-mitochondria contact sites in co-overexpression and endogenous conditions we used Duolink-Proximity Ligation Assay (PLA) coupled with staining of mitochondria (MitoTracker) and confocal microscopy. PLA signals corresponding to ORP5-ORP8 interaction were observed throughout the cell in both endogenous and co-overexpression (HA-ORP5 and 3XFLAG-ORP8) conditions (Fig. 2a). The specificity of this assay and of the antibodies used was confirmed by the strong decrease in PLA signals for endogenous ORP5-ORP8 interaction in cells with ORP5 and ORP8 knocked down (Fig. 2b, S3a). Likewise, a significant increase in ORP5-8 PLA signals was induced by the overexpression of these proteins (Fig. 2b) Close association of PLA spots to mitochondria, indicating localization at ER-mitochondria contact sites, was measured after segmentation of the mitochondrial network by Imaris (Fig. 2a, right panel; S3b). Interestingly, the majority of ORP5-8 PLA signals localized at ER-mitochondria contact sites (52% of endogenous ORP5-8 and 50% of co-overexpressed ORP5-8) (Fig. 2a, 2c). The localization of ORP5-8 PLA signals to ER-PM contact sites was analyzed in HeLa cells transfected with RFP-PH-PLCδ, to stain the PM, and Mito-BFP, to label mitochondria. However, only a minor pool of ORP5-8 PLA spots was found in contact with the PM (4% of endogenous ORP5-8 and 5% of co-overexpressed ORP5-8) (Fig. 2d-f). The localization of ORP5-ORP8 PLA spots to the ER, including MAMs, was confirmed in cells co-expressing the ER protein Sec22b and the mitochondrial-targeted Mito-BFP (Fig. S3c).

**Figure 2.**
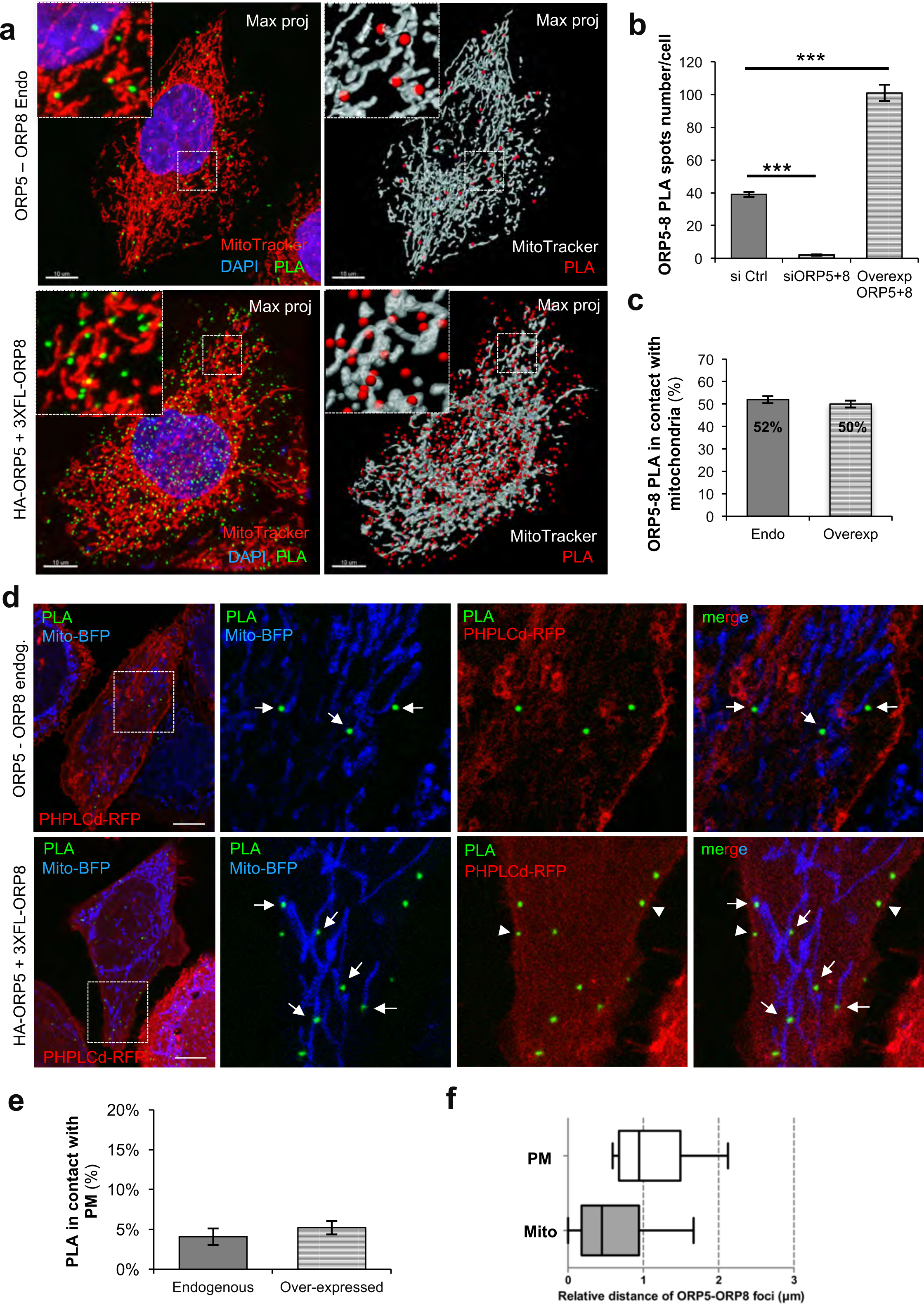
The main localization of the endogenous ORP5-8 complex is ER-mitochondria contact sites. (a) Representative confocal images of ORP5-ORP8 interaction in HeLa cells detected by Duolink PLA (green spots) in endogenous (ORP5-ORP8 Endo) and overexpressing (HA-ORP5 + 3xFL-ORP8) conditions, and their respective 3D representation by Imaris. Images are presented as maximum projection of all layers. Insets show magnifications of the boxed regions. Scale bar, 10 μm. (b) Quantification of the number of ORP5-ORP8 PLA interactions in Control (siCtrl, n=39 cells), ORP5 and ORP8 knockdown (siORP5+8, n=38 cells), and in overexpression of ORP5 and ORP8 (Ovrexp ORP5+8, n=35 cells), showing that the downregulation or the upregulation of both ORP5 and ORP8, respectively, reduces and increases the number of interaction stablished between these two proteins. Statistical analysis: P values were determined by unpaired student’s t-test, ***P<0,001. (c) Quantification of ORP5-ORP8 PLA interaction localized to ER-Mitochondria contact sites in control (Endo, n= 33) and HeLa cells overexpressing (HA-ORP5 + 3xFL-ORP8, n=27 cells) showing that about 50% of ORP5-ORP8 interactions occurs at MAM. (d) Representative confocal images of ORP5-ORP8 PLA interaction (green spots) detected in HeLa cells overexpressing PHPLCd-RFP, Mito-BFP (ORP5-ORP8 Endo), or in HeLa cells overexpressing PHPLCd- RFP, Mito-BFP, ORP5 and ORP8 (HA-ORP5 + 3xFL-ORP8). Images are presented as individual layers. Scale bar, 10 μm. (e) Quantification of ORP5-ORP8 PLA signal localized to ER-plasma membrane contact sites indicate that about 5% of the total ORP5-ORP8 interactions occurs to these subdomains of the ER membranes, in both control (Endogenous, n=6 cells) and HeLa cells overexpressing ORP5 and ORP8 (Overexpressed, n=14 cells). (f) Box plot of ORP5-ORP8 endogenous PLA spots distance (in μm) to mitochondria and plasma membrane (box around median value, whiskers 10%- 90%) evidencing that the majority of ORP5-ORP8 interactions were detected in a close proximity to mitochondria (<0.38 μm) and distant from the plasma membrane (≥0.38 μm).

Overall these data reveal for the first time that the main sites where ORP5 and ORP8 localize and interact in endogenous physiological conditions are the ER-mitochondria contact sites, and not the ER-PM contacts.

### ORP5/8 physically interact with the mitochondrial intermembrane space bridging (MIB) complex facing cristae junctions

To investigate whether ORP5/8 localize to specific ER-mitochondria contact subdomains we performed a morphological analysis of ORP5 localization by immuno- EM (IEM) on ultrathin cryosections from HeLa cells transfected with HA-ORP5 or EGFP-ORP5 (as endogenous ORP5/8 levels are too low to be detected by IEM). We previously reported that about 20% of ORP5 or ORP8 gold particles were associated to ER-mitochondria contact sites when individually expressed (Galmes et al., 2016). The advantage of analyzing ORP5 localization is its preferential localization to contact sites (20% at ER-mitochondria contacts and 60% ER-PM contacts), when expressed alone, as compared to ORP8, which remains also largely present within the reticular ER (60% of ORP8 vs 20% of ORP5) (Galmes et al., 2016). Interestingly, the majority of ORP5 gold particles at ER-mitochondria contact sites was found to localize to ER elements in a very close proximity (86% within 0-100 nm distances, 50% of which within 50 nm) to the CJ (arrow, Fig. 3a-b), tubular structures that connect the IMM to the cristae. To exclude that ORP5 localization near CJ is not a consequence of its distribution throughout the ER membranes, we sought to determine if other ER proteins have a similar frequency of proximity to CJ. Thus, we compared ORP5 localization to Sec61β, an ER protein present in ER elements widely distributed throughout the cells and very little at ER-mitochondria contacts (Galmes et al., 2016). Co-immunolabeling of EGFP-ORP5 or EGFP-Sec61β and the luminal ER protein disulfide isomerase (PDI) confirmed ORP5 localization to ER elements close to CJ (arrow, Fig. 3c) but not of Sec61β, the bulk of which localized on ER membranes distant from the CJ (0% within 0-100nm and 69% >200nm distance) even when close to mitochondria (Fig. 3b and arrowheads in Fig. 3c). To determine whether other contact sites proteins would be similarly enriched at CJ we analyzed the localization of PTPIP51, a mitochondrial tether known to localize at ER-mitochondria contact sites (Stoica, De Vos et al., 2014) and also a binding partner of ORP5/8 (Galmes et al., 2016). A small pool of PTPIP51- HA could be detected near CJ where they colocalized with EGFP-ORP8 or EGFP- ORP5. However, the majority of PTPIP51 randomly distributed throughout the mitochondrial surface (Fig. 3d-e). Hence, our results support the conclusion that ORP5/8 specifically localize to ER-mitochondria contact sites closely associated to CJ. Interestingly, in yeast, CJ were shown to be closely associated to OMM-IMM contact sites tethered by the MICOS complex (Harner et al., 2011). IEM analysis using Mic60- EGFP, an EGFP-tagged construct of the human orthologue of the central component of the MICOS complex, confirmed that human Mic60, similarly to its yeast orthologue, preferentially localizes to the IMM in close proximity of CJ and in the cristae that arise from them (arrow, Fig. 3a). These results suggest that ER-mitochondria contact sites where ORP5/8 localize could be physically connected to the intra-mitochondrial membrane contact sites near CJ.

**Figure 3.**
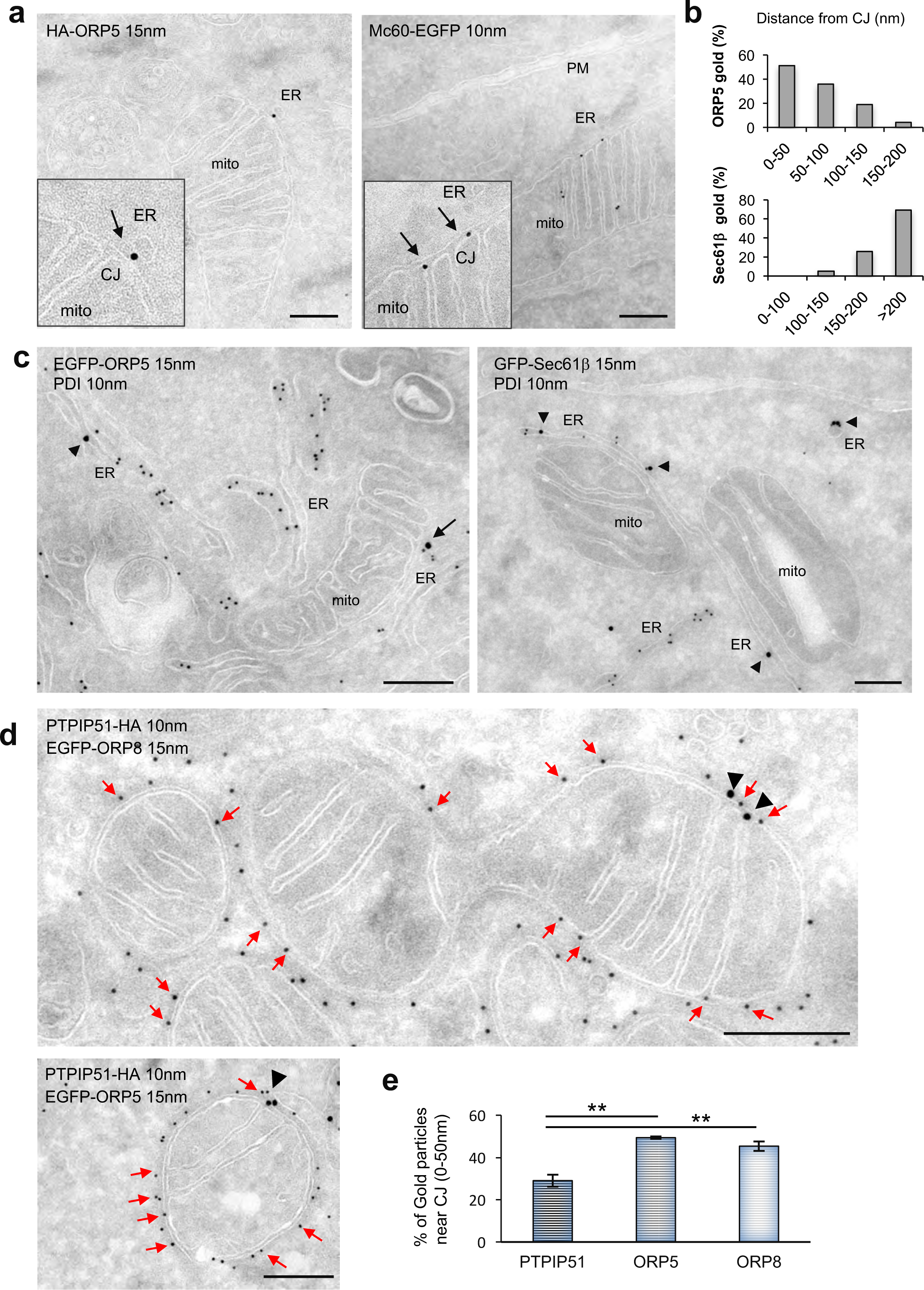
ORP5 localizes at ER-mitochondria contact sites near cristae junctions (CJ). (a) Electron micrographs of ultrathin cryosections of HeLa cells transfected with HA-ORP5 or Mic60-EGFP and immunogold stained with anti-HA or anti-GFP (10 or 15 nm gold), showing ORP5 localization at ER-mitochondria contacts in close proximity to CJ (arrow) and the localization of the MICOS complex (Mic60) at CJ (arrows). Scale bar 250 nm. (b) Quantification of the proximity of HA-ORP5 and EGFP-Sec61b gold particles to the CJ. Results are presented as the percentage of ORP5 or Sec61b gold particles at specific ranges of distance (in nm) from CJ. 150 gold particles were counted on randomly selected cell profiles in each sample. (c) Electron micrographs of ultrathin cryosections of HeLa cells transfected with EGFP-ORP5 or GFP-Sec61b and immunogold labeled with anti-GFP (15 nm gold) and anti-PDI (10nm gold). Note ORP5 localization at ER-mitochondria contacts near CJ (arrow) and Sec61b localization to ER membranes not in contact with the mitochondria membranes (arrowheads). Scale bar 250 nm. (d) Electron micrographs of ultrathin cryosections of HeLa cells transfected with EGFP-ORP8 or EGFP-ORP5 and PTPIP51-HA and immunogold labeled with anti-GFP (15 nm gold) and anti-HA (10nm gold). Note ORP8 and ORP5 specific localization at ER-mitochondria contacts near CJ (black arrowheads) and PTPIP51 localization to all the the mitochondria surface including CJ (red arrows). Scale bars 250 nm. (e) Quantification of PTPIP51-HA, EGFP-ORP8 and EGFP-ORP5 proximities to CJ. Results are presented as the % of gold particles within 0-50nm distance from CJ±SEM. 286 gold particles for PTPIP51 and 150 gold particles for ORP5 and ORP8 were counted on randomly selected cell profiles in each sample. Statistical analysis: unpaired student’s t-test, **P<0.01.

To identify binding partners of ORP5/8 at ER-mitochondria contact sites we carried out a MS-analysis on GFP-pull downs from cells expressing EGFP-ORP5, EGFP- ORP5ΔPH (an ORP5 variant lacking the PM-targeting PH domain that is localized at ER-mitochondria but not at ER-PM contact sites) or EGFP alone as a control (Fig. 4a, S4a). As expected, the highest hit detected in both EGFP-ORP5 and EGFP-ORP5ΔPH pull-downs was ORP8. In accord to our previous study (Galmes et al., 2016), the mitochondrial protein PTPIP51 was also detected in the mass spectrometry analysis. Interestingly, several new outer mitochondrial membrane proteins (listed in Fig. 4a) were also found as major hits. Among these proteins, the MIB component SAM50 and the MICOS central subunit Mic60, binding partner of SAM50 at OMM-IMM contact sites (Ott et al., 2015, Tang, Zhang et al., 2020), had the highest scores (Fig. 4a). Interestingly, SAM50 and Mic60 showed a higher interaction score in EGFP-ORP5ΔPH immunoprecipitates, as compared to EGFP-ORP5. WB analysis using anti-actin as loading control showed that EGFP-ORP5ΔPH overexpression did not alter the amount of SAM50 and Mic60 proteins as compared to the overexpression of neither EGFP-ORP5 nor EGFP alone (Fig. S4d), indicating that the higher interaction scores of these proteins in the EGFP-ORP5ΔPH immunoprecipitates were not due to their increased levels. Of note, metaxin-2 was also detected in the MS of immunoprecipitated EGFP-ORP5 and EGFP-ORP5ΔPH, although its score was lower than SAM50 and Mic60 and in the EGFP-ORP5 was below the assigned threshold (50) (Fig. 4a).

**Figure 4.**
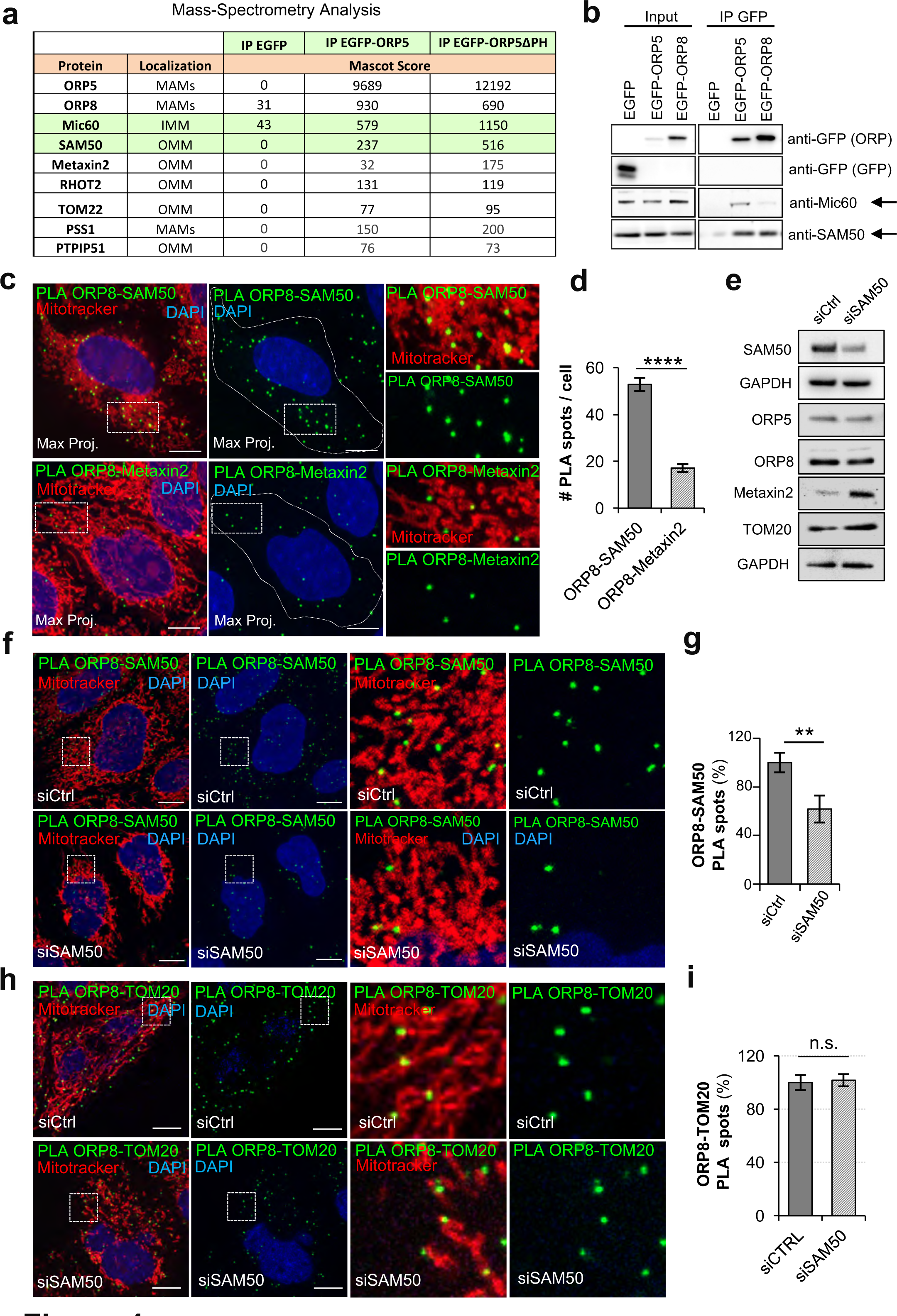
ORP5 and ORP8 interact with the MIB/MICOS complex at ER- mitochondria contacts. (a) Identification of mitochondrial proteins associated to mitochondrial outer or inner membranes (OMM, IMM) that interact with EGFP-tagged ORP5 constructs by mass spectrometry. Note the presence of some proteins of the MIB complex: Mic60, SAM50 and metaxin2 and of their interacting partner RHOT2. Interaction scores (Mascot scores) of Mic60, SAM50 and metaxin2 with the EGFP-ORP5ΔPH construct are stronger than with EGFP-ORP5. (b) Western blot analysis of ORP5, ORP8, SAM50 and Mic60 in immuno-precipitated samples obtained from lysates of HeLa cells transfected with EGFP-ORP5, EGFP-ORP8 or EGFP alone. ORP5 and ORP8 were detected using antibodies against GFP. (c) Representative confocal images showing endogenous ORP8-SAM50 and ORP8-metaxin2 PLA interactions (green), mitochondrial network (MitoTracker, red) and nuclei (DAPI, blue) in HeLa cells. Images are displayed as maximum projection of all layers. Scale bar, 10 μm. (d) Quantitative analysis of endogenous ORP8-SAM50 and ORP8-metaxin2 PLA interactions in HeLa cells. Data are shown as mean values ±SEM of n= 36 cells, ORP8-SAM50; n= 39 cells, ORP8-Metaxin2. Statistical analysis was performed using unpaired student’s t-test, with ****, P<0.0001. (e) Western blot analysis of ORP5, ORP8, SAM50, TOM20 and metaxin2 in protein lysates obtained from HeLa cells treated with scrambled siRNA (siCtrl) and from Hela cells treated with siRNA targeting SAM50 (siSAM50). (f) Confocal images of control (siCtrl) and SAM50 (siSAM50) knockdown HeLa cells showing endogenous interaction of ORP8-SAM50 by Duolink PLA (green) at MAMs. Mitochondria are labeled by MitoTracker (red) and nuclei by DAPI (blue). Images are presented as maximum projection of all layers. Insets show magnifications of the boxed regions. Scale bar, 10 μm. (g) Quantification of endogenous ORP8-SAM50 PLA signals in Control and SAM50 knockdown HeLa cells, showing the reduction of about 50% of ORP8-SAM50 PLA in SAM50 knockdown cells as compared to control. Bars indicate mean values ±SEM. Number of cell analysed: siCtrl (n=30), siSAM50 (n=22). Statistical analysis: P values were determined by unpaired student’s t-test, **P<0.01. (h) Representative confocal images showing endogenous ORP8-TOM20 PLA interactions (green), mitochondria (MitoTracker, red) and nuclei (DAPI, blue) in Control and SAM50 knockdown HeLa cells. Images are displayed as single layers. Scale bar, 10 μm. (i) Quantitative analysis of endogenous ORP8-TOM20 PLA interactions in HeLa cells. Data are shown as mean values ±SEM of n=49 cells, siCtrl; n= 49 cells, siSAM50, n.s; not significant.

To confirm ORP5/8 interaction with SAM50 and Mic60, GFP-pull down experiments from HeLa cells expressing EGFP-ORP5, EGFP-ORP8 or EGFP alone were carried out (Fig. 4b). Consistent with the MS data, endogenous SAM50 and Mic60 were recovered with both EGFP-ORP5 and EGFP-ORP8 but not with EGFP alone, confirming specific biochemical co-purification of ORP5 and ORP8 with SAM50 and Mic60.

Next, to determine the domains involved in the interaction of ORP5/8 with the MIB/MICOS complex, GFP-pull down experiments were carried out from cells expressing EGFP-tagged ORP5 (EGFP-ORP5ΔPH) or ORP8 (EGFP-ORP8ΔPH) PH domain deleted constructs (Fig. S4a-b), and compared to the full-length proteins (EGFP-ORP5 and EGFP-ORP8) or to the EGFP alone. In accord with the MS data, the deletion of the PH domain increased ORP5 and ORP8 interaction with SAM50, as compared to the full-length proteins (Fig. S4b). Confocal analysis of ORP5 and SAM50 localization in cells expressing EGFP-ORP5 or EGFP-ORP5ΔPH and stained with an anti-SAM50 antibody confirmed the stronger enrichment of the ΔPH ORP5 construct at ER elements in contact with the SAM50-labeled mitochondria as compared to the full-length ORP5 (Fig. S4c).

As the PH domain is not required for the interaction with SAM50, we further investigated the role of the other domains of ORP5 in such interaction by immunoprecipitating ORP5 deletion mutants for the ORD or the TM domains (EGFP- ORP5ΔORD, EGFP-ORP5ΔTM) (Fig. S4a-b). While the deletion of the ORD domain did not affect the interaction between ORP5 and SAM50, the deletion of the TM domain decreased the amount of SAM50 co-immunoprecipitated with ORP5, indicating that ORP5 should be properly anchored to the ER to localize at ER-mitochondria contact sites to interact with SAM50 (Fig. S4b).

To confirm the interaction between ORP5/8 and SAM50 at endogenous level we took advantage of the available antibodies against ORP8 and SAM50 from different species and analyzed their interaction using PLA (duolink) by confocal imaging in HeLa cells. PLA signals corresponding to ORP8-SAM50 endogenous interaction were detected at ER-mitochondria contact sites in control cells (Fig. 4c). Interestingly, PLA signals were also detected for ORP8-metaxin 2, although the number of spots was significantly minor (Fig. 4c-d), correlating to the lower MS score of metaxin 2 (Fig. 4a). Then, to further verify the specificity of ORP8-SAM50 PLA we analyzed their interaction in cells where SAM50 was downregulated by RNAi (siSAM50) (4e-f). A decrease of about 40% of ORP8-SAM50 PLA was found in siSAM50 cells (Fig. 4f-g), in accord with the decrease of the levels of SAM50 protein of about 40-50% assessed by WB (Fig. 4e), and validating the specificity of the ORP8-SAM50 interaction. Similarly, the KD of SAM50 decreased also the PLA interaction between ORP8 (and ORP5) and metaxin 2 (Fig. S5a-d). To further investigate if this effect was specific for protein components of the MIB complex we performed PLA interaction assays of ORP8 (and ORP5) with another outer mitochondrial membrane protein TOM20, that is not a bona-fide component of this complex. Interestingly, TOM20 also interacted with ORP8 and, to a minor extent with ORP5, but these interactions were not decreased by the KD of SAM50 (Fig. 4h-i, S5e-f). Also, these effects were not due to decreased levels of ORP5, ORP8, metaxin 2 and TOM20 proteins, as assessed by WB (Fig. 4e). Together, our data reveal the specificity of the effects of SAM50 KD on ORP5/8 reciprocal interaction and with components of the MIB complex.

To investigate a possible role of SAM50 and Mic60 in regulating the levels of ORP5/8 at MAMs we analyzed their endogenous interaction by PLA and confocal microscopy in HeLa cells where either SAM50 or Mic60 were knocked-down by siRNA. Interestingly, a significant decrease (of about 40%) in ORP5/8 interaction in close proximity to mitochondria (corresponding to MAM) was found in both SAM50 and Mic60 KD cells, as compared to control cells (Fig. 5a-b), indicating a synergistic effect of SAM50/Mic60 on the ORP5-ORP8 interaction at MAM. Further localization analysis of endogenous ORP5 and ORP8 in siSAM50 cells confirmed decreased levels of these proteins at MAM (Fig. 5 c-f).

**Figure 5.**
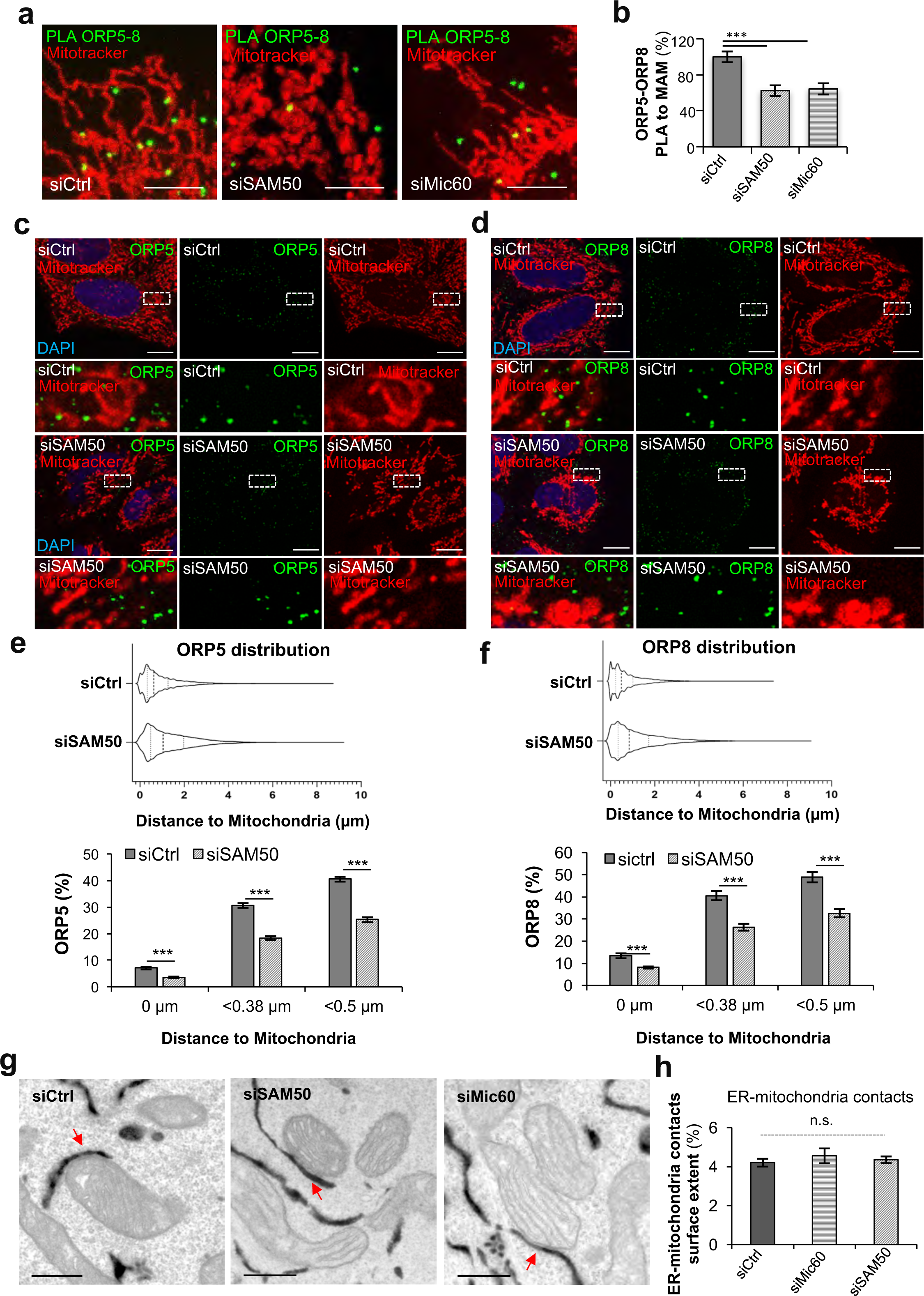
SAM50 and Mic60 knockdowns induce a decrease in ORP5-8 interaction at MAMs but do not alter the abundance of ER-mitochondria contact sites. (a) Confocal images of a region of Ctrl (siCtrl), SAM50 (siSAM50) and Mic60 (siMic60) knockdown HeLa cells showing endogenous interaction of ORP5-8 by Duolink PLA (green) near mitochondria (MitoTracker, red). Scale bar, 5 μm. (b) Quantification of ORP5-8 PLA signals in Control, SAM50 and Mic60 knockdown HeLa cells, showing the decrease of about 40% of ORP5-ORP8 PLA in SAM50 and Mic60 knockdown cells as compared to control. Bars indicate mean values ±SEM. Number of cell analysed: siCtrl (n=33), siSAM50 (n=19), siMic60 (n=24). Statistical analysis: unpaired sudent’s t-test. ***P<0.001. (c-d) Confocal images showing endogenous ORP5 (green) and ORP8 (green) localization in relation to mitochondria (MitoTracker, red) in Ctrl (siCtrl) and SAM50 (siSAM50) knockdown HeLa cells. Nuclei are stained with DAPI (blue). Images are represented as single layers. Magnifications of wite boxed regions are shown for each condition. Scale bar, 10 μm. (e-f) Quantitative analysis of ORP5 and ORP8 distribution in relation to their distance to mitochondria in HeLa cells treated with scrambled siRNA (siCtrl) and siRNA targeting SAM50 (siSAM50). Violin plots display overall frequency and distribution of ORP5 and ORP8 labelling in relation to their distance to mitochondria. Median values (wider dotted line) anti-ORP5; siCtrl= 0.61 µm, siSAM50= 1.04 µm; and anti-ORP8: siCtrl= 0.48 µm, siSAM50= 0.84 µm. Interquartile range (narrow dotted lines) anti-ORP5; siCtrl 25% and 75% percentile= 0.32 and 1.26 µm, siSAM50 25% and 75% percentile= 0.48 and 1.97 µm. Anti-ORP8: siCtrl 25% and 75 % percentile= 0.23 and 1.01 µm, siSAM50 25% and 75% percentile= 0.34 µm and 1.7 µm. Column charts display the % of ORP5 and ORP8 labelling that distance from mitochondria 0 µm, <0.38 µm and <0.5 µm. Data are expressed as mean values ±SEM. Number of cell analyzed: anti-ORP5: siCtrl n=37, siSAM50 n= 40; anti-ORP8: siCtrl n=22, siSAM50 n= 29. Statistical analysis: P values were determined by unpaired student’s t-test, *** P<0.001. (g) Representative electron micrographs of HeLa cells treated with Ctrl siRNAs or siRNAs against SAM50 or Mic60 and transfected with HRP-KDEL. Red arrows indicate ER-mitochondria contact sites. Scale bar, 500 nm. (h) Quantifications of the extent of ER-mitochondria contact sites in Ctrl, Mic60 and SAM50 knockdown cells expressing HRP-KDEL. Data are shown as % of the ER in contact with mitochondria (mitochondria occupancy) ±SEM, n = 30 for siCtrl, n = 20 cell profiles for siMic60 and siSAM50 and 1000 mitochondria; n.s; not significant.

To verify the possibility of an indirect effect of SAM50 or Mic60 silencing on the morphology and abundance of MAMs we carried out an ultrastructural analysis by conventional EM and HRP-KDEL EM (carrying a horseradish peroxidase (HRP) tagged with an ER retention motif to stain the ER) in Mic60 or SAM50 silenced cells. Morphological analysis by conventional EM showed that transient KD of SAM50 or Mic60 induce formation of multilamellar cristae, almost devoid of CJ (Fig. S6a-b), complementing previous observations by other groups through stable disruption of the MICOS/MIB functions (Ding, Wu et al., 2015, Ott et al., 2015). However, in both SAM50 and Mic60 KD cells ER-mitochondria contact sites were still present and their morphology not altered (Fig. 5g, S6a). Quantitative morphological analysis by HRP-KDEL EM in control and SAM50 or Mic60 silenced cells confirmed that the abundance of ER-mitochondria contact sites was not altered by SAM50 or Mic60 KD (Fig. 5h), indicating that the effects on ORP5/8 interaction at MAMs were not indirect due to a global rearrangement of the ER-mitochondria contact sites.

Overall our data reveal a novel interaction between ORP5/8 and the MIB/MICOS complex at ER subdomains associated with intra-mitochondrial membrane contacts facing CJ, and a direct role of the MIB/MICOS complex in ORP5/8 targeting/interactions at MAMs.

### ORP5/8 and the MIB/MICOS complex regulate PS-to-PE conversion at the ER-mitochondria interface

The role of ORP5 and ORP8 in lipid transport at ER-mitochondria contacts still remains to be established. We tested whether ORP5 and ORP8 could mediate PS transport at the ER-mitochondria interface by measuring levels of mitochondrial PE in the mitochondrial fraction isolated from HeLa cells (as in (Galmes et al., 2016)) where ORP5 or ORP8 were transiently silenced by RNAi. As the ER-derived PS is the major precursor for mitochondrial PE, if ORP5 and ORP8 mediate non-vesicular transport of PS from the ER to the mitochondria, their absence should lead to a reduction of mitochondrial PE. We chose to use a transient KD as it overcomes the limits and/or compensatory effects on lipid transport/biosynthetic pathways that other stable approaches could induce. The purity of mitochondria and of the other subcellular fractions was verified in control, ORP5 and ORP8 KD conditions by WB by probing the samples for cytochrome c as mitochondrial marker and IP3R-3 as a MAM-enriched marker (Fig. 6a). All markers were highly enriched in their respective fractions and were absent in the others, and, as previously shown (Galmes et al., 2016), ORP5 and ORP8 were enriched in the MAM fraction and absent in the mitochondria fraction of control cells (Fig. 6a). On the contrary, they were strongly suppressed in ORP5 and ORP8 KD cell lysates and in the respective MAM fractions (Fig. 6a). MS- lipidomic analysis revealed a specific reduction of PE levels in mitochondria isolated from ORP5 and ORP8 KD cells, of 34% and 20%, respectively, as compared to control cells (Fig. 6b), while PE levels from total cells were unchanged. Additionally, the mitochondrial levels of two other phospholipids (PI and PC) were not perturbed by ORP5/8 depletion (Fig. S8a). Additionally, the decrease in PE could be due to a decrease in the protein levels of the PS-decarboxylase (PISD) or the PS-Synthase 1 (PSS1) enzymes mediating PS-to-PE conversion on mitochondria or PS synthesis in the ER, respectively. To exclude this possibility, we analyzed the protein levels of PISD and PSS1 by WB in ORP5, ORP8, ORP5+ORP8 KD cells and compared them to control cells (Fig. S8b). We found no significant difference but rather a slight increase in the enzymes upon ORP5 and/or ORP8 KDs. These data suggest that ORP5 and ORP8 could transfer PS at ER-mitochondria contact sites.

**Figure 6.**
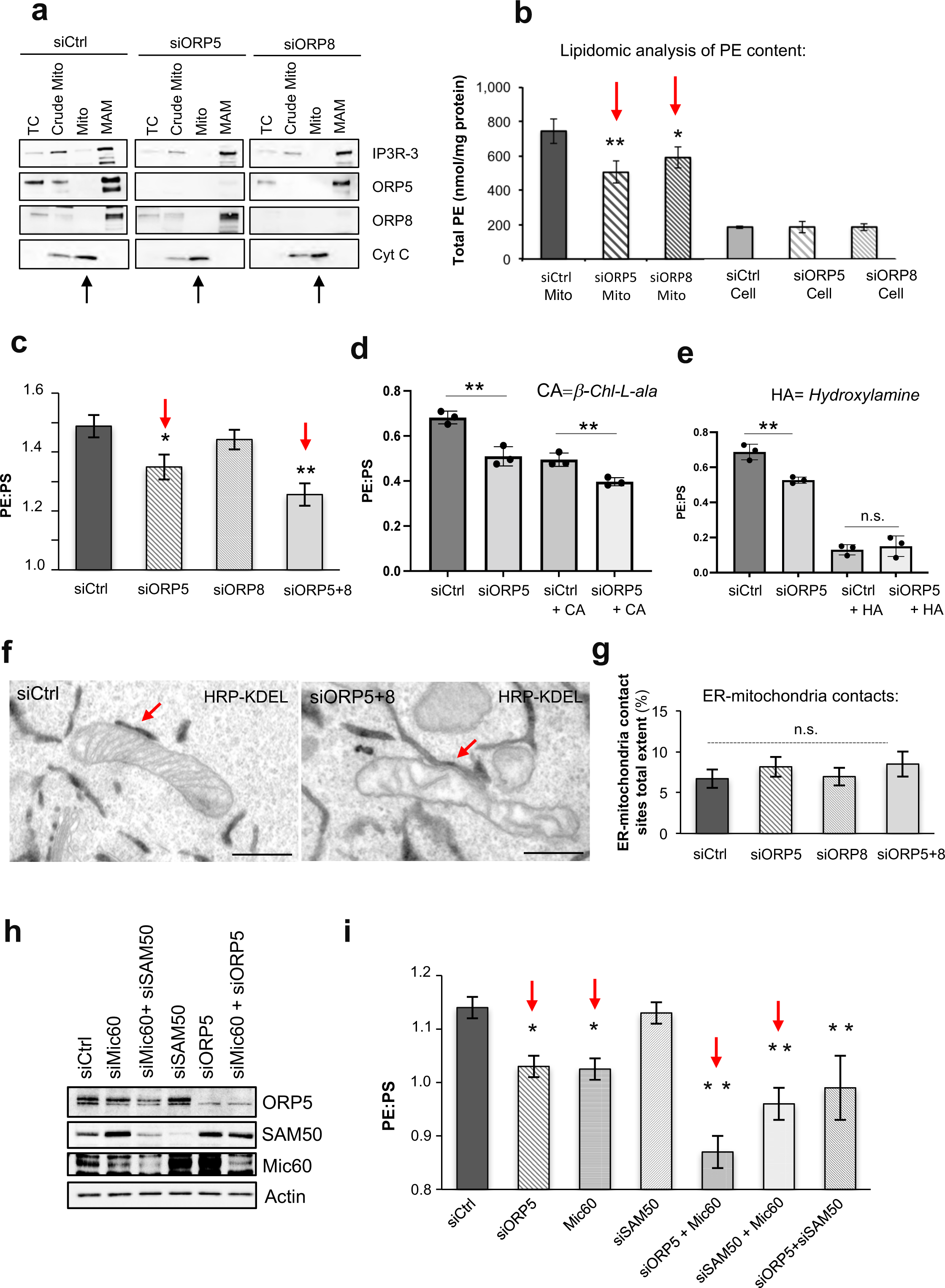
ORP5/8 and the MIB/MICOS complex regulate levels of PS-derived mitochondrial PE. (a) Crude mitochondria, mitochondria, and MAM fractions were purified from Ctrl, ORP5 and ORP8 siRNA-treated HeLa cells. Equal amounts of protein from each fraction were loaded on a 4–20% gradient SDS–PAGE gel and immunoblotted using anti-ORP5, anti-ORP8, anti-IP3R-3 (MAM protein), and anti-cytochrome c (mitochondrial protein). Mito, mitochondria; MAM, mitochondria-associated ER membrane. (b) Mass spectrometry (MS)-based quantification of the PE content (nmol/mg protein) of mitochondria isolated from Ctrl, ORP5 or ORP8 knockdown cells and of Ctrl, ORP5 or ORP8 knockdown intact cells. Data are shown as mean of three independent replicates ±SEM. Statistical analysis: unpaired student’s t-test, *P<0.05, **P<0.01. (c) HeLa cells transfected with siCtrl, siORP5, siORP8 or siORP5+ORP8 RNAi oligos were incubated with L-[^3^H(G)]serine (30.9 Ci/mmol) for 18 hours. After extraction and separation of lipids by TLC, PS and PE spots were scraped and analyzed for [^3^H] radioactivity, as described under “Methods”. Each condition was analyzed in triplicate in each of three independent biological replicates. Data are presented as mean of PE:PS ratio ±SEM. Statistical analysis: unpaired student’s t-test, *P<0.05, **P<0.01 compared to Ctrl. (d) Cells transfected with siCtrl and siORP5 oligos were treated with 1 mM ý-Chloro-L-alanine (CA, ý-Chl-L-ala, inhibitor of Ser-palmitoyltransferase) or untreated, then pulsed with 7 μCi/ml of [3H(G)]serine for 1 hour and chased for 4 hours in serum-free DMEM + ý-Chloro-L- alanine, before analysis. n.s. not significant, **P<0.01 compared to Ctrl. n.s. not significant. (e) Similar experiments were performed in presence of 5 mM hydroxylamine (HA, inhibitor of phosphatidylserine decarboxylase). **P<0.01, n.s. not significant. (f) Electron micrographs of HRP-KDEL-expressing HeLa cells treated with Ctrl siRNAs (siCtrl) or siRNAs against ORP5 and ORP8 (siORP5+siORP8). Red arrows indicate ER- mitochondria contact sites. Scale bar, 500 nm. (g) Quantifications of the total extent of ER- mitochondria contact sites in siCtrl, siORP5, siORP8 and siORP5+8 cells expressing HRP-KDEL. Data are shown as % of the ER surface lenght in contact with mitochondria ±SEM, n = 20 cell profiles and ±900 mitochondria; n.s; not significant. (h) Western analysis showing ORP5, SAM50, Mic60 and Actin levels in protein lysates from HeLa cells treated with siRNA against Ctrl, ORP5, Mic60 or SAM50. Arrow indicates the specific band for Mic60. (i) Radiometric measurement of PS-to-PE conversion in the indicated siRNAs. Data are presented as mean of PE:PS ratio ±SEM. Each condition was performed in triplicate in each of the independent biological replicates (n = 5 for siCtrl and siORP5; n = 4 for siSAM50; n = 3 for the other siRNAs conditions). Statistical analysis: unpaired student’s t-test , *P<0.05, **P<0.01 compared to Ctrl.

The ORD domain of ORP8 (ORD8) has been shown to transfer PS between liposome membranes in counter-transport with PI4P or PIP_2_ *in vitro* (Chung et al. 2015; Ghai et al, 2017). However, the lipid transfer activity of ORP5 ORD domain (ORD5) has not been studied so far. Also, the ability of ORD5 and ORD8 to transfer PS independently of a gradient of PI4P or PIP_2_ or other phospholipids than PS has never been addressed. Thus, we purified the recombinant ORD5 and ORD8 from bacteria (Fig. S7a) and compared their ability to transport fluorescent phospholipids (TopFluor-PS, -PC or -PE) from donor to acceptor liposomes *in vitro*. Donor liposomes containing fluorescent phospholipids and biotinylated lipids were first immobilized on streptavidin beads and then mixed with acceptor liposomes in the presence or absence of ORP5/8 ORD domains (Fig. S7b). After 1 hour at 37°C, acceptor liposomes were recovered from the supernatant and their fluorescence was measured (Fig. S7b). Our results show that both ORD5 and ORD8 transfer PS, but not PC and PE, from donor to acceptor liposomes (Fig. S7c). They also reveal that ORD5 and ORD8 share equivalent ability to transfer PS *in vitro*. To confirm that fluorescent lipids were indeed transferred to the acceptor liposomes, a fraction of the reaction supernatant was floated on a Nycodenz density gradient by ultracentrifugation and the fluorescence in the top fraction of the gradient (containing floated acceptor liposomes) was measured (Fig. S7d). Fluorescence of TopFluor-PS in the acceptor liposomes was maintained after their floatation, confirming its effective transfer between liposomes *in vitro*.

In subsequent experiments, we measured the levels of mitochondrial PE newly synthesized from the ER-derived PS by using a radiometric PS-to-PE conversion assay *in situ* (Shiao, Lupo et al., 1995) in silenced or control HeLa cells (Fig. 6c). This assay allows the monitoring of PS transfer from the ER to mitochondria by measuring the levels of radioactive PS and PS-derived PE by thin layer chromatography (TLC) after incorporation of radioactive L-[^3^H(G)]-serine into the cells. A significant decrease in the levels of newly synthetized PE was found in ORP5 and in ORP5+ORP8 KD cells (Fig. 6c). The decrease was stronger in ORP5+ORP8 KD cells, indicating a cooperative effect of ORP5 and ORP8 in this process. A slight, not statistically significant decrease was also found in ORP8 KD cells, suggesting a major role of ORP5 compared to ORP8 in PS transfer at ER-mitochondria contact sites *in situ*. As [^3^H]-serine radioactivity could be incorporated to PE also *via* an alternative pathway involving sphingosine (Hanada, Nishijima et al., 1992), we next sought to address the contribution of this pathway to PE labeling, by repeating the experiments in control and ORP5 KD cells in the presence throughout pulse and chase of [3H]-serine of β-chloro-L-alanine, an inhibitor of serine palmitoyltransferase (Chen, Born et al., 1993). Although a decrease in newly synthesized PE was observed in siCtrl cells upon addition of β-chloro-L-alanine, PS-to-PE conversion was significantly reduced in β- chloro-L-alanine treated and untreated ORP5 knockdown cells (∼17% and ∼23% respectively), as compared to control (Fig. 6d). We then verified that the fraction of PE decreased upon ORP5 KD was indeed derived from conversion of PS by PISD by measuring PS-to-PE conversion in the presence of hydroxylamine, an inhibitor of PISD activity (Fig. 6e). Treatment with 5 mM hydroxylamine robustly inhibited the PS to PE conversion in both control and ORP5 KD cells, the level of residual conversion being similar in both of these cells, further confirming that the reduction in mitochondrial PE induced by depletion of ORP5 is essentially due to the decrease in PS transfer from the ER to the mitochondria (Fig. S8e). The β-chloro-L-alanine and hydroxylamine data also suggest that at least 80% of the serine labeling of PE occurs via the phosphatidylserine decarboxylase reaction in HeLa cells, in accord with a previous work in another cell type, the Baby Hamster Kidney fibroblasts (BHK cells), showing that only certain minor PE species are labeled from the sphingosine-PE pathway (Heikinheimo & Somerharju, 1998).

To test whether the effects on PS transport at ER-mitochondria contacts were specific for ORP5 and ORP8 loss of function or simply due to a decrease of ER-mitochondria contacts induced by their KD, we quantified the abundance of ER-mitochondria contact sites by EM in control and ORP5, ORP8, and ORP5+ORP8 KD cells. To facilitate the visualization of the ER we transfected the cells with a HRP-KDEL construct that stains the ER with a dark signal. Our quantifications revealed that ORP5, ORP8 or ORP5+ORP8 KDs did not affect the extent of ER-mitochondria contacts (Fig. 6f-g). The total mitochondrial mass (quantified as mitochondrial surface/cell) was also unchanged (Fig. S8c). These results indicate that ORP5 and ORP8 act as LTPs rather than tethers (Fig. 7, S8e). Even a modest reduction (22-27%) of mitochondrial PE levels in mammalian cells has been shown to profoundly alter the morphology of mitochondrial cristae as well as mitochondria functions (Tasseva, Bai et al., 2013). Accordingly, 52% of mitochondria in ORP5+ORP8 double-KD cells display aberrant cristae morphology versus 9% in control cells (Fig. 6f, S8d). These defects in cristae morphology were also observed by conventional EM (Fig. S6a) and were similar to those previously shown in the case of ORP5 and ORP8 individual KD (Galmes et al., 2016). Interestingly, the % of mitochondria with altered morphology in ORP5+ORP8 double-KD cells was higher as compared to ORP8 KD (Galmes et al., 2016), reflecting the stronger effect of ORP5+ORP8 double-KD on PS transport at ER-mitochondria contact sites (Fig. 6c, S8d).

**Figure 7.**
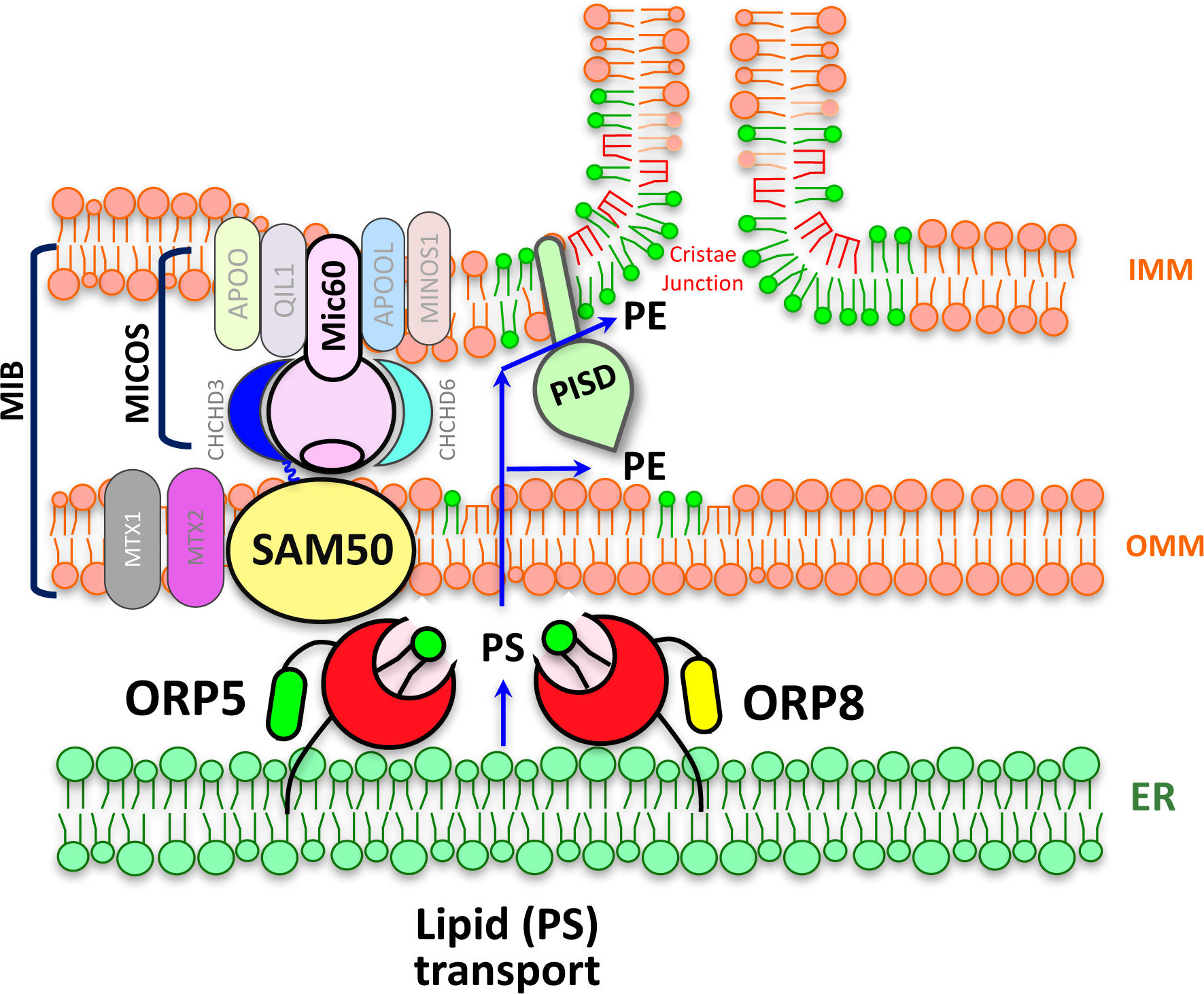
PS transport at ER-mitochondria contact site subdomains associated to MIB/MICOS complex. ORP5/8 mediate the transfer of PS from ER to mitochondria at ER-mitochondria membrane contact sites. This transfer occurs at ER subdomains facing the cristae junctions (CJ) where ORP5/8 interacts with SAM50 and Mic60, key proteins of the MIB complex. This interaction facilitates the transfer of PS from ER to the mitochondrial membranes at the level of CJ and PS conversion into PE, a phospholipid that plays a critical role in cristae organization and mitochondrial function.

Decreased levels of PE strongly affect the organization of the mitochondrial respiratory supercomplexes (Tasseva et al., 2013). We had previously shown that ORP5 KD induces a reduction in the basal mitochondrial oxygen consumption rate (OCR_BAS_), indicative of reduced respiratory activity (Galmes et al., 2016). However, it remains still questioned whether ORP8 could also reduce OCR_BAS_ and if this reduction could be exacerbated under metabolic stress conditions. Thus, we monitored mitochondrial OCR in control, in ORP5 and in ORP8 KD cells in basal and in stress conditions (Fig. S9). ORP5 KD induced a significant reduction in both OCR_BAS_ (∼37%) and OCR upon FCCP treatment (OCR_FCCP_) (∼36%). Interestingly, also ORP8 KD induced a significant decrease in OCR_BAS_ (∼31%) and OCR_FCCP_ (∼29%), although the decreases were less prominent than upon ORP5 KD. These data uncover a novel role of ORP5 and ORP8 in preserving mitochondrial respiratory activity in basal and in stress conditions, and a major impact of ORP5 in this process, in accord with its major role in PS transfer at ER-mitochondria contact sites.

To test whether the interaction of ORP5 with the MIB complex could facilitate the non-vesicular transfer of PS from the ER to the mitochondrial membranes (and consequently synthesis of mitochondrial PE) we performed radiometric PS-to-PE conversion assay in cells depleted of ORP5, SAM50 or Mic60 alone or in combination by RNAi. Robust KD of ORP5, SAM50 or Mic60 was confirmed by western blotting after 48 hours (Fig. 6h). Analysis of PS-derived newly synthetized PE revealed a significant decrease in PE in ORP5 and Mic60 KD cells (Fig. 6i). Moreover, the double-KD of ORP5 and Mic60 had an additive effect, supporting a cooperation of these two proteins in the regulation of PE synthesis. However, the levels of PE were not changed in SAM50 KD cells as compared to control. This could be explained by the fact that other subunits of the MIB complex might compensate for its depletion. Indeed, protein levels of Mic60 and metaxin 2 were strongly increased upon SAM50 KD (Fig. 4e, 6h). Interestingly, the double-silencing of SAM50 and either ORP5 or Mic60 had a significant impact on PE synthesis that was even stronger as compared to the individual KDs (Fig. 6i).

EM analysis of mitochondria morphology in all KD conditions revealed that downregulation of ORP5, SAM50 and Mic60, individually or in combination, induces formation of multilamellar cristae, almost devoid of CJ (Fig. S6a-b). However, in all KD conditions ER-mitochondria contact sites were still present and their morphology was not altered (Fig. 5g-h, Fig S6a), indicating that the effects on PS-derived PE synthesis were specifically due to ORP5, Mic60 or SAM50 loss of function effects on PS transport at ER-mitochondria contacts or on the maintenance of intra-mitochondrial membrane bridges, respectively.

Overall our results reveal that ORP5/8 cooperate with the MIB/MICOS complex to regulate the transfer of PS from the ER to the mitochondrial membranes necessary for synthesis of mitochondrial PE and consequently for maintaining mitochondrial cristae morphology and respiration (Fig. 7).

## DISCUSSION

In this study, by using a combination of biochemical and imaging approaches, we uncover for the first time the endogenous localization of ORP5 and ORP8, revealing that they are mainly localized at ER-mitochondria contact sites and providing novel evidence for a physiological relevance of the ORP5/8 complex at MAMs (Fig. 1-2 and Fig. S1-3). So far, ORP5 and ORP8 localization have been only studied in conditions where one of these two partner proteins were expressed in high excess as compared to the other one. Previous studies, including one from our group, have shown that overexpression of ORP5 induces an increase of ER-PM contacts where the protein also localizes (Chung et al., 2015, Galmes et al., 2016). Consequently, several following studies have addressed the role of ORP5 and ORP8 at ER-PM contacts, overlooking their function at MAM. However, the increase in cortical ER observed upon ORP5 overexpression does not reflect the physiological abundance of ER-PM contacts, as the cortical ER in non-specialized cells generally covers not more than 5% of the plasma membrane surface. Indeed, we have found that the increased localization of ORP5 to cortical ER when over-expressed alone does not reflect its endogenous localization when it is in complex with ORP8, which is instead enriched at MAMs. Our findings have important implications for a better understanding of the physiological localization and function of ORP proteins, but also of other proteins that assemble in multimeric complexes at ER-mediated membrane contact sites, such as the E-Syts (Giordano, Saheki et al., 2013).

Our study reveals that ORP5/8 physically interact with SAM50 and Mic60, two key components of the MIB/MICOS complex that anchor the IMM to the OMM at the level of CJ (Huynen et al., 2016, Ott et al., 2015, Wollweber et al., 2017). The biochemical interaction between ORP5 and SAM50/ Mic60 uncovers the existence of a novel physical link between ER-mitochondria contact sites involved in lipid transport and intra- mitochondrial membrane contacts. ORP5 localization by IEM at ER-mitochondria contact sites near the CJ, where Mic60 and MICOS complex also reside, further confirms the existence of such tripartite membrane contact site structure. Moreover, KD of SAM50 and Mic60 does not affect ER-mitochondria contact sites, but specifically perturbs ORP5/8 interactions with themselves and with metaxin 2, another component of the MIB complex at MAMs.

Importantly, here we describe a new function of ORP5 and ORP8 in the maintenance of mitochondrial levels of PE, an essential phospholipid of mitochondria, providing the first evidence of mammalian LTPs directly mediating non-vesicular transfer of PS (lipid precursor of mitochondrial PE) from the MAMs to the mitochondria at ER-mitochondria contact sites.

Other LTPs have been recently identified at ER-mitochondria contact sites in mammalian cells. Hirabayashi et al. recently showed that the SMP-containing protein PDZD8 is involved in ER-mitochondria tethering and in the regulation of Ca^2+^ dynamics in mammalian neurons (Hirabayashi et al., 2017). Although PDZD8 is a structural and functional paralogous of the Mmm1 subunit of the ERMES complex (Wideman et al., 2018), its function in lipid transport at ER-mitochondria contact sites remains unclear. Recently, the mammalian LTP VPS13A has been shown to localize to contact sites including ER-mitochondria contacts (Kumar et al., 2018). VPS13A contains a lipid-binding domain (VPS13α) that has the ability to harbor multiple phospholipids at once and transfer them between liposomes *in vitro*. However, its role in lipid transfer at ER-mitochondria membrane contact sites *in situ* has not yet been established. Differently from the SMP and VPS13α domains that can simultaneously host multiple phospholipids, the ORD domain of Osh/ORPs forms a cavity that can host only one lipid at a time (Maeda, Anand et al., 2013, Wang, Ma et al., 2019).

An important finding of our work is that ORP5 and ORP8 KD do not affect the extent of ER-mitochondria contact sites, revealing that the main function of ORP5/8 at MAMs is lipid transfer and not membrane tethering. This is a unique feature among the LTPs that have been identified so far at MAMs. For instance, KD of PDZD8 and VPS13A result in a decrease of ER-mitochondria contact sites (Hirabayashi et al., 2017, Kornmann, Currie et al., 2009, Kumar et al., 2018), making it difficult to dissect the lipid transfer activity from the tethering function of these LTPs. Thus, ORP5/8 represent a so far a unique example of LTPs that specifically mediate lipid transport at ER-mitochondria contact sites, independently of membrane tethering, and that can be used as unique tools to specifically study lipid transport at MAMs.

ORP5 and ORP8 have been previously shown to counter-exchange PS with the PM phosphoinositides PI4P and PIP_2_ at ER-PM contact sites in HeLa cells (Chung et al., 2015, Ghai, Du et al., 2017). However, PI4P and PIP_2_ are not present on the mitochondrial membranes, while PE is highly abundant in these membranes, in addition to being an essential lipid of all biological membranes. Our *in vitro* data show that the ORD domains of ORP5 and ORP8 transport PS, but not other phospholipids such as PE and PC, indicating a specific role of ORP5/8 in PS transport and excluding the possibility that ORP5/8 might also participate in the transport of a fraction of PE back to the ER. It is possible that ORP5/8 cooperate with other LTPs, such as VPS13A, for the exchange of other lipids (including PE) at ER-mitochondria contact sites. Importantly, we have confirmed the role of ORP5/8 in PS transfer by measuring a specific decrease of PS-derived mitochondrial PE in ORP5 depleted HeLa cells *in situ* (and even more upon ORP5+8 silencing) as well as a reduction of total PE in mitochondria isolated from these cells (Fig. 6a-e, S8a-b). Accordingly, ORP5/8 KD affects cristae morphology and the respiratory function of mitochondria (Galmes et al., 2016)(Fig. 6f-g, S6, S8d), all phenotypes that are expected in the case of even a mild decrease in mitochondrial PE (Joshi et al., 2012, Steenbergen et al., 2005);(Tasseva et al., 2013). Our data also suggest that the gradient of PS at ER-mitochondria contacts is sufficient to trigger the ORP5/8-mediated transport of PS from the MAMs, where it is highly enriched, to the mitochondria membranes, where it is rapidly converted into PE and is therefore present at a very low concentration. Our findings have important implications in the general field of LTPs, as they suggest that the same LTP can use different means to transfer lipids depending on the local gradients present at the specific membrane contact sites where it is localized.

Importantly, we also show that the *de-novo* synthesis of mitochondrial PE requires both ORP5/8 at ER-mitochondria contact sites and the MIB/MICOS complex at intra- mitochondrial OMM-IMM contact sites (Fig. 6i). Interestingly, recent evidence in yeast suggests that, in addition to the classical PE synthesis at the IMM by the IMM-localized PS-decarboxylase PISD, PE can also be synthesized *in trans* on the OMM (Aaltonen, Friedman et al., 2016). Thus, it is possible that this alternative pathway, which requires MIB/MICOS tethering function to bring the mitochondrial intermembrane domain of PISD close to the OMM for synthesis of PE, is conserved also in mammalian cells. The cooperation of ORP5 (through its lipid transfer activity at ER-mitochondria contact sites) with SAM50 and Mic60 (through their OMM-IMM tethering function) could facilitate the movement of PS from the ER to the IMM where it is converted to PE, through the classical PE synthesis pathway or through PISD function *in trans* on the OMM. Taken together these findings provide the first evidence of a physical and functional link between ER-mitochondria contacts and intra-mitochondrial membrane contacts maintained by the MIB/MICOS complexes, to facilitate transport of PS from the ER to the mitochondria and PE synthesis on the mitochondrial membranes (Fig. 7).

In conclusion, our data reveal that: 1) ORP5/8 form a protein complex that is endogenously enriched at ER-mitochondria contacts; 2) ORP5/8 constitute a molecular machinery mediating PS transfer at ER-mitochondria contact sites but not ER-mitochondria tethering; 3) ER-mitochondria contacts where ORP5/8 localize are physically associated with intra-mitochondrial contacts, maintained by the MIB/MICOS complex, to facilitate the transport of PS from the ER to mitochondria membranes. Overall our study provides a first molecular clue on how lipids are transported at ER-mitochondria contact sites and novel functional insight into the complex interplay of the ER with the mitochondria and the intra-mitochondrial membrane contacts and associated machinery.

## STAR METHODS

### RESOURCES AVAILABILITY

#### Lead contact

Further information and requests for resources and reagents should be directed to and will be fulfilled by the lead contact, Francesca Giordano. All unique reagents generated in this study are available from the Lead Contact.

### Materials availability

All unique reagents generated in this study are available from the Lead Contact.

## EXPERIMENTAL MODEL AND SUBJECT DETAILS

### Cell line

HeLa cells were used for KD, transfection, biochemistry, light and electron microscopy, and lipidomic studies. HeLa cells were cultured in Dulbecco’s Modified Eagle Medium (DMEM, Life Technologies) supplemented with 10% FBS (Life Technologies) and 1% penicillin/streptomycin (P/S, 100 units/ml penicillin and 10 μg/mL streptomycin, Life Technologies) and maintained at 37°C with 5% CO_2_.

## METHOD DETAILS

### cDNA plasmids and molecular cloning

EGFP-ORP5, EGFP-ORP8, EGFP-ORP5ΔPH, EGFP-ORP8ΔPH, EGFP-ORP5ΔORD and EGFP-ORP5ΔTM were described in (Galmes et al., 2016). RFP- Sec22b (Gallo, Danglot et al., 2020). The following reagents were kind gifts: GFP- Sec61β from T. Rapoport (Harward University)(Shibata, Voss et al., 2008), PHPLC8- RFP (Chung et al., 2015); Mito-BFP (Addgene: # 49151); ssHRP-KDEL from T. Schikorski (Schikorski, Young et al., 2007); GST-ORD8 (ORD ORP8, corresponding to aa 328-767) from P. De Camilli (Chung et al., 2015).

Cloning of HA-ORP5, Mic60-GFP and GST- ORD5 (ORD ORP5) : cDNAs of ORP5 (full-length), Mitoflin (full-length from FLAG- Mic60 (Ott et al., 2015) and GST- ORD5 (corresponding to aa 265-703), were amplified by PCR. In all PCR reactions, Herculase II fusion DNA polymerase (Agilent) was used.

Primers used were (coding sequence shown in lowercase): 5’ AgeI-HA-ORP5_Fw GGCGGC *ACCGGT* cgccacc ATGTACCCATACGATGTTCCAGATTACGCT atgaaggaggaggccttcctc 3’ XhoI-STOP-ORP5_Rv GGC *CTCGAG* ctatttgaggatgtggttaatg 5’ KpnI- Mic60_Fw AGACCCAAGCTT *GGTACC* atg

3’ BamHI-GC- Mic60_Rv GTAATC *GGATTC* GC ctctggct

5’ SalI-TC-ORD5_Fw GCACAG GTCGAC TC gagacccctggggccccggt

3’ NotI-STOP-ORD5_Rv GCACA *GCGGCCGC* ctactgtggccggagggctggtcg For the HA-ORP5 cloning the PCR product (carrying the HA tag at the N-terminus of ORP5) was ligated between AgeI and XhoI in the pEGFP-C1 vector (Clontech) and replacing the GFP- with the HA-tag. For the other clonings the PCR products were ligated between KpnI and BamHI for Mic60, and between SalI and

NotI for ORD5, in the pEGFP-N1 vector (Clontech) to generate Mic60-EGFP or in the pGEX-6P-1 to generate GST-ORD5.

### Cell transfection

HeLa cells were seeded in 13mm glass coverslips (Agar Scientific) or in 10 cm dishes and were cells were transfected with 1 μg or 24 ug DNA, respectively, using Lipofectamine2000 (Invitrogen) according to the manufacturer’s instruction. Cells were cultured for 24 hours prior to analysis.

### RNA interference

ORP5, ORP8, SAM50 and Mic60 KD in Hela cells was achieved by siRNA transfection. HeLa cells were seeded in 13 mm coverslips (Agar Scientific) or in 10 cm dishes transfected with 200 nM siRNA using Oligofectamine (Invitrogen) following the manufacturer’s instruction. Cells were cultured for 48 hours prior to analysis.

Double-stranded siRNAs were derived from the following references:

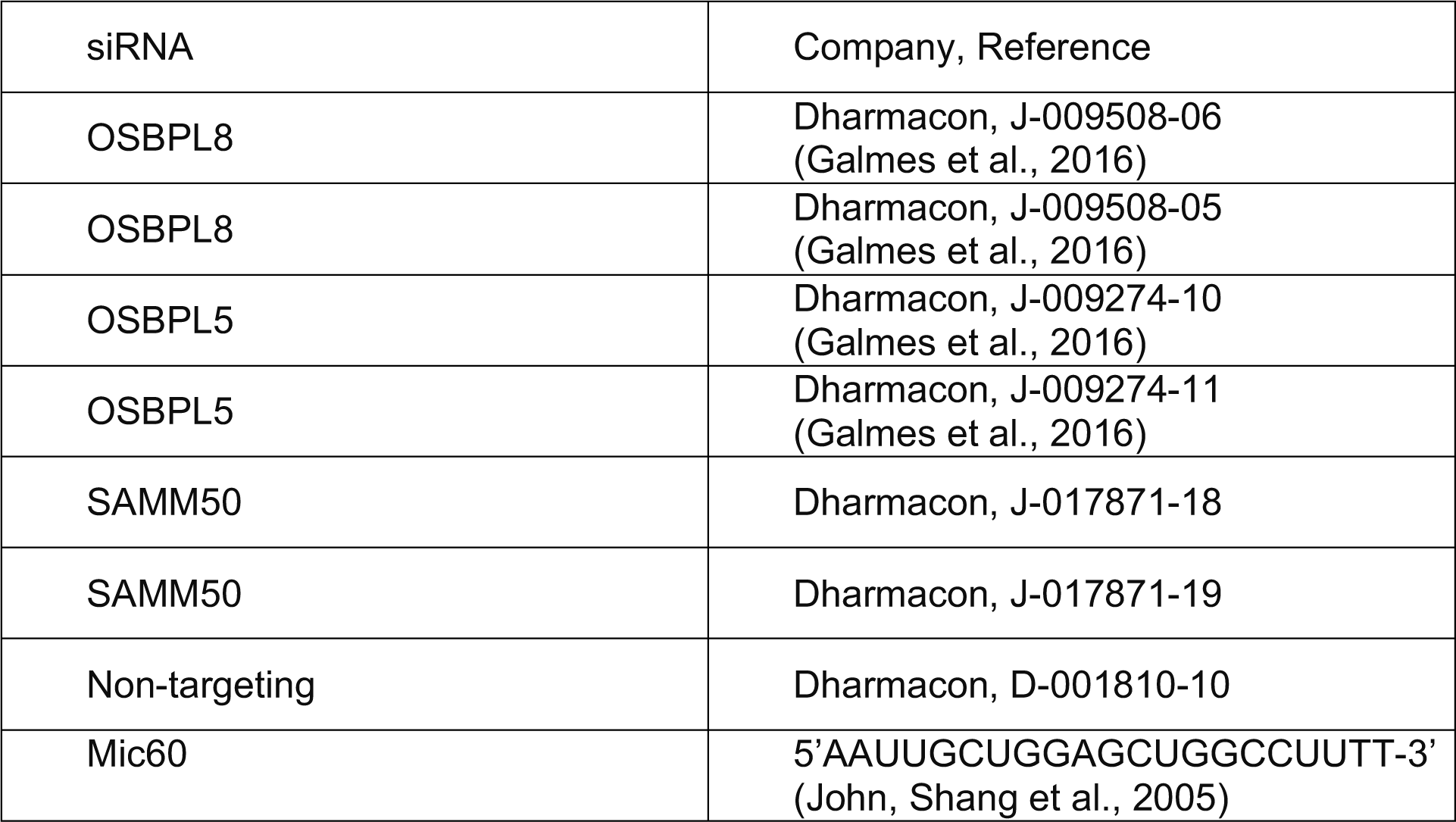

### mRNA analyses by quantitative reverse transcriptase PCR (qPCR)

Total RNA was isolated from HeLa cells transfected with siRNAs for 48 hours as described above, by using a Purelink™ kit (Ambion/Thermo Scientific, Foster City, CA). The RNA (0.5 μg per specimen) was reverse transcribed with a SuperScript VILO™ cDNA synthesis kit (Invitrogen/Thermo Scientific, Carlsbad, CA) according to the manufacturer’s protocol. Quantification of the mRNAs of interest was carried out on a Roche Lightcycler™ 480 II instrument by using SYBR-Green I Master mix (Roche, Basel, Switzerland) and primers specified in the Table bellow. Succinate dehydrogenase complex. subunit A, the mRNA of which remained markedly stable under the present conditions, was employed as a reference housekeeping mRNA. Relative mRNA levels were calculated by using the –ýýCt method.

Sequences of the primers used for qPCR:

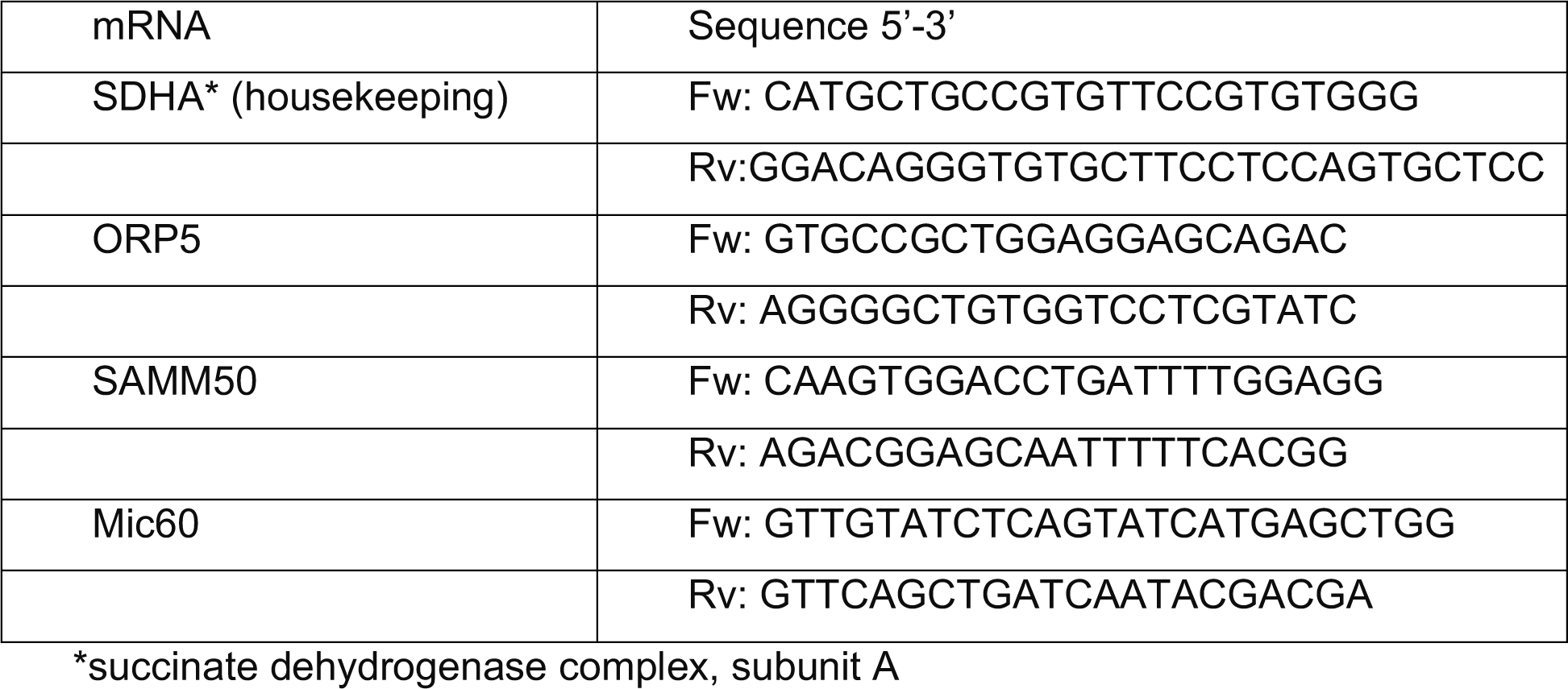

### Immunoprecipitation of ORPs

HeLa cells transfected with EGFP-tagged ORPs were washed in cold PBS and lysed on ice in lysis buffer [50 mM Tris, 120 mM NaCl, 40 mM Hepes, 0.5% digitonin, 0.5% CHAPS, pH 7.36, and protease inhibitor cocktail (Roche)]. Cell lysates were then centrifuged at 21 000 g for 20 min at 4°C. Supernatants were then incubated with Chromotek GFP-trap agarose beads (Allele Biotech) for 1 hour at 4°C under rotation. Subsequently beads were washed in 0.1 M phosphate buffer. After extensive washes in cold lysis buffer, immunoprecipitated proteins bound to the beads were processed for Mass Spectrometry analysis or incubated in sample buffer (containing 2% SDS) and then boiled for 1 min at 97 °C. In the latter case immunoprecipitates were loaded and separated in 10% SDS–PAGE gel and immunoblotting was carried out.

### Western blotting

For immunoblotting, cells were resuspended in lysis buffer [50 mM Tris, 150 mM NaCl, 1% Triton X-100, 10 mM EDTA, pH 7.2, and protease inhibitor cocktail (Roche)]. Cell lysates were then centrifuged at 21 000 g for 20 min at 4°C. The supernatants were boiled in reducing SDS sample buffer. Proteins isolated from HeLa cells or obtained by immunoprecipitation were were subjected to SDS-PAGE gels for electrophoresis separation. The separated proteins were transferred to 0.45 μm Nitrocellulose membrane (GE Healthcare). The membrane was blocked by 5% non-fat milk in TBST buffer (TBS buffer with 0.1% Tween 20) for 1 h at room temperature and washed 3 times, for 5 min each, with TBST. Then the membrane was incubated with the primary antibodies (antibodies and dilutions listed below) at 4°C overnight. The membrane was washed 3 times wash with TBST incubated with the peroxidase-conjugated anti-rabbit IgG secondary antibody (1:5000, in 5%milk in TBST) or anti-mouse IgG secondary (1:5000, in 5%milk in TBST) at room temperature for 1h, followed with washing and detection using the enhanced chemiluminescence (ECL) detection kit (Cytiva). For Western blot quantification, bands of protein of interest were detected using ChemiDoc™ Imaging Systems (Life Science Research, Bio-Rad) and analyzed using Image Lab™ Software. All data are presented as mean ± standard error of the mean (SEM) of three experimental replicates.

| Primary Antibodies (Company & Reference) | Dilution |
| --- | --- |
| IP3R-3 (BD Transduction Laboratories, 610312) | 1:2000 |
| ORP5 (SIGMA, HPA038712) | 1:1000 |
| ORP5 (SIGMA, HPA058727) | 1:1000 |
| β-Actin (Abcam, ab8226) | 1:1000 |
| ORP8 (GeneTex, GTX121273) | 1:1000 |
| SAM50 (SIGMA, HPA034537) | 1:1000 |
| Mic60 (Proteintech, 10179-1-AP) | 1:1000 |
| metaxin2 (Santa Cruz, sc-514231) | 1:1000 |
| TOM20 (BD Transduction Laboratories, 612278) | 1:10000 |
| PDI (GeneTex, GTX30716) | 1:500 |
| VAPB (SIGMA, HPA013144) | 1:500 |
| PSS1 (PTDSS1, SIGMA, HPA016852) | 1:300 |
| PISD (SIGMA, HPA031090) | 1:300 |
| GFP (Roche, 11814460001) | 1:1000 |
| Secondary Antibodies (Company & Reference) | Dilution |
| anti-mouse-HRP (GE Healthcare, NA931V) | 1:10000 |
| anti-rabbit-HRP (GE Healthcare, NA934V) | 1:10000 |

### Mass spectrometry-proteomic analysis

Mass Spectrometry (MS) analysis was carried out by the proteomics/mass spectrometry platform in IJM (http://www.ijm.fr/plateformes/spectrometrie-de-masse). Briefly, after washes with binding buffer, immunoprecipitations beads were rinsed with 100 μl of NH4HCO3 25 mmol/l. Proteins on beads were digested overnight at 37°C by sequencing grade trypsin (12.5 μg/ml; Promega Madison, Wi, USA) in 20 μl of NH4HCO3 25 mmol/l. Digests were analysed by an Orbitrap Fusion (Thermo Fisher Scientific, San Jose, CA) equipped with a Thermo Scientific EASY-Spray nanoelectrospray ion source and coupled to an Easy nano-LC Proxeon 1000 system (Thermo Fisher Scientific, San Jose, CA). MS/MS data were processed with Proteome Discoverer 1.4 software (Thermo Fisher scientific, San Jose, CA) coupled to an in house Mascot search server (Matrix Science, Boston, MA; version 2.4.2). MS/MS datas were searched against SwissProt databases with Homo sapiens taxonomy. The Mascot score for a protein is the summed score for the individual peptides, e.g. peptide masses and peptide fragment ion masses, for all peptides matching a given protein. For a positive protein identification, the mascot score has to be above the 95% confidence level. In Mascot, the ions score for an MS/MS match is based on the calculated probability, P, that the observed match between the experimental data and the database sequence is a random event. The reported score is -10Log_10_(P). A score of 200 indicates a probability of 10^-20^. Scores greater than 70 are significant, while scores lower than 40 should not be considered or carefully validated at MS/MS level (source: http://www.matrixscience.com/help/interpretation_help.html). We thus set up a Mascot score threshold of 50.

### Cell fractionation

HeLa cells (100x10^6^ cells) were harvested 48 hours after transfection with siRNA oligos and washed with PBS by centrifugation at 600 g for 5 min. The cell pellet was resuspended in starting buffer (225 mM mannitol, 75 mM sucrose and 30 mM Tris-HCl pH 7.4) and homogenized using a Tissue Grinder Dura-Grind®, Stainless Steel, Dounce (Wheaton). The homogenate was centrifuged three times at 600 g for 5 min to remove nuclei and unbroken cells. The crude mitochondria was pelleted by centrifugation at 10 000 g for 10 min. To separate MAM and pure mitochondria fractions, the pellet was resuspended in MRB buffer (250 mM mannitol, 5 mM HEPES and 0.5 mM EGTA, pH 7.4) and layered on top of different concentrations of Percoll gradient (225 mM mannitol, 25 mM HEPES, 1 mM EGTA pH 7.4 and 30% or 15% Percoll). After centrifugation at 95 000 g for 30 min, two dense bands containing either the pure mitochondria or MAM fraction were recovered and washed twice with MRB buffer by centrifugation at 6300 g for 10 min to remove residual Percoll and residual contamination. MAM was pelleted by centrifugation at 100 000 g for 1 hour. MAMs and pure mitochondria pellets were resuspended in Lysis Buffer (50 mM Tris, 150 mM NaCl, 1% Triton X-100, 10 mM EDTA, pH 7.2, and protease inhibitor cocktail) and protein concentrations were determined by Bradford assay. Equal amount of proteins were loaded on 4-20% gradient SDS-PAGE gels (Biorad) and immunoblotting was carried out. Pure mitochondria were processed for MS-lipidomic analysis.

### Mass Spectrometry-lipidomic analysis

700 μl of homogenized cells were mixed with 800 μl 1 N HCl:CH_3_OH 1:8 (v/v), 900 μl CHCl_3_ and 200 μg/ml of the antioxidant 2,6-di-tert-butyl-4-methylphenol (BHT; Sigma Aldrich). The organic fraction was evaporated using a Savant Speedvac spd111v (Thermo Fisher Scientific). Lipid pellets were reconstituted in running solution (CH_3_OH:CHCl_3_:NH_4_OH; 90:10:1.25; v/v/v). Phospholipid species were analyzed by electrospray ionization tandem mass spectrometry (ESI-MS/MS) on a hybrid triple quadrupole/linear ion trap mass spectrometer (4000 QTRAP system; Applied Biosystems SCIEX) equipped with a TriVersa NanoMate (Advion Biosciences) robotic nanosource. Phospholipid profiling was executed by (positive or negative) precursor ion or neutral loss scanning at a collision energy of 35 eV for neutral loss 141 (phosphatidylethanolamine (PE)). Phospholipid quantification was performed by multiple reaction monitoring (MRM), the transitions being based on the neutral losses or the typical product ions as described above. The MRM dwell time was set to 100 ms and typically the signal was averaged over 20 cycles. Lipid standards used were PE25:0 and PE43:6 (Avanti Polar Lipids). The data were corrected for isotope effects as described by (Liebisch, Lieser et al., 2004).

### Immunofluorescence

HeLa cells were seeded on 13 mm glass bottom coverslips (Agar Scientific). KD or/and transfected cells were incubated with 1 μM MitoTracker (mitochondrial marker, Invitrogen) for 30 min at 37°C, 5% CO_2_ and then fixed with 4% PFA/PBS for 15 min at room temperature. Fixed cells were then washed in PBS and incubated with 50 mM NH4Cl/PBS for 15 min at room temperature. After washing with PBS and blocking buffer (1% BSA/ 0.1% Saponin in PBS), cells were incubated with primary antibodies diluted in blocking buffer for 1 hour at room temperature and then with fluorescently-labeled secondary antibodies. After washing with blocking buffer and then PBS, coverslips were mounted on microscopy slides and images were acquired on Confocal inverted microscope SP8-X (DMI 6000 Leica). Optical sections were acquired with a Plan Apo 63x oil immersion objective (N.A. 1.4, Leica) using the LAS-X software. Fluorescence was excited using either a 405nm laser diode or a white light laser, and later collected after adjusting the spectral windows with GaAsP PMTs or Hybrid detectors. Images from a mid-focal plane are shown. Images were processed and fluorescence was analysed off line using Image J. For co-localization analysis of fluorescent signals, the acquired images were processed using the JACoP plugin in ImageJ to assess the Pearson’s correlation coefficient. The obtained values, ranging from 0 to 1 (1=max correlation), indicated the association between the signals analysed. To assess the distance between ORP5 or ORP8 and mitochondria confocal images were processed by Imaris, using a similar approach described in the PLA section.

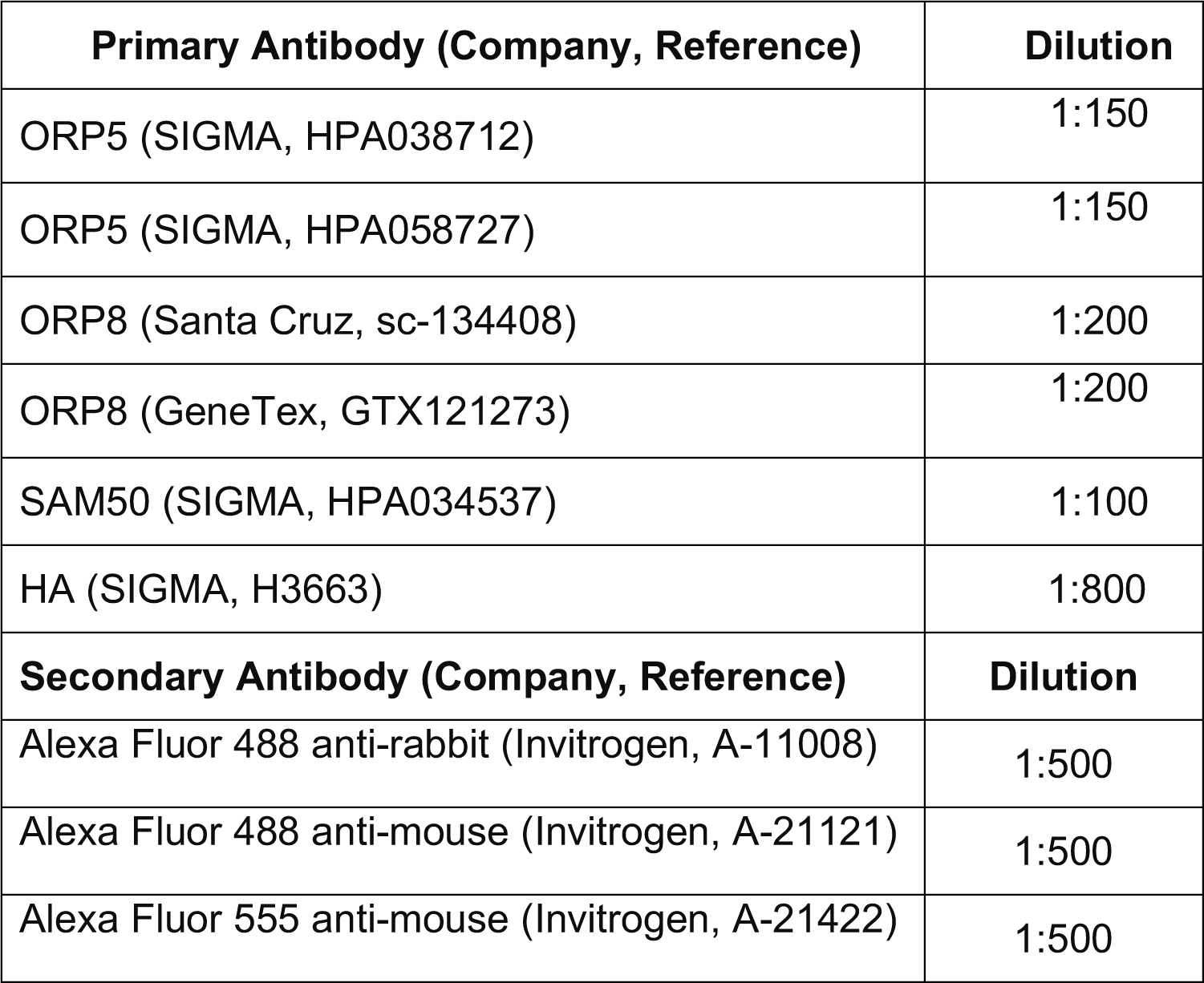

| Primary Antibody (Company, Reference) | Dilution |
| --- | --- |
| ORP5 (SIGMA, HPA038712) | 1:150 |
| ORP5 (SIGMA, HPA058727) | 1:150 |
| ORP8 (Santa Cruz, sc-134408) | 1:200 |
| ORP8 (GeneTex, GTX121273) | 1:200 |
| SAM50 (SIGMA, HPA034537) | 1:100 |
| HA (SIGMA, H3663) | 1:800 |
| Secondary Antibody (Company, Reference) | Dilution |
| Alexa Fluor 488 anti-rabbit (Invitrogen, A-11008) | 1:500 |
| Alexa Fluor 488 anti-mouse (Invitrogen, A-21121) | 1:500 |
| Alexa Fluor 555 anti-mouse (Invitrogen, A-21422) | 1:500 |

### *In situ* proximity ligation assay (PLA)

The protein-protein interactions in fixed HeLa cells were assessed using *in situ* PLA (Duolink^®^SIGMA) according with the manufacturer’s instructions. Briefly, HeLa cells seeded on 13 mm glass coverslips (Agar Scientific) were incubated with MitoTracker Red (mitochondrial marker, Invitrogen, 1 μM in DMEM) for 30 min at 37°C or co-transfected with PHPLCd-RFP (plasma membrane marker) plus Mito-BFP (mitochondrial marker). Cells were thereafter fixed with 4% PFA for 30 minutes at room temperature and incubated with primary antibodies (dilutions in the table below) in blocking solution (1% BSA, w/v 0.01% saponin, w/v, in PBS) for 1h at room temperature. PLUS and MINUS PLA probes (anti-murine and anti-rabbit IgG antibodies conjugated with oligonucleotides, 1:5 in blocking solution) were then incubated with the samples for 1h at 37°C. Coverslips were thereafter washed in 1x wash buffer A and incubated with ligation solution (5x Duolink^®^ Ligation buffer 1:5, ligase 1:40 in high purity water) for 30 min at 37°C. After the ligation step, cell samples were washed in 1x wash buffer A and incubated with the polymerase solution (5x Amplification buffer 1:5, polymerase 1:80 in high purity water) for 1h40min at 37°C. Polymerase solution was washed out from the coverslips with 1x wash buffer B and 0.01x wash buffer B. Vectashield Mounting Medium with or without DAPI (Vector Laboratories) was used for mounting. Images were acquired on Confocal inverted microscope SP8-X (DMI 6000 Leica). Optical sections were acquired with a Plan Apo 63x oil immersion objective (N.A. 1.4, Leica) using the LAS-X software. Fluorescence was excited using either a 405nm laser diode or a white light laser, and later collected after adjusting the spectral windows with GaAsP PMTs or Hybrid detectors. Images from a mid-focal plane or maximal projection of all layers are shown. Images were processed and the number and the distance of PLA dots to mitochondria and to the plasma membrane were assessed using the Imaris software (v 9.3, Bitplane). Briefly, segmented 3D images (PLA foci identified as “spots”, mitochondria identified as “surfaces”, and plasma membrane represented as “cell” were generated from confocal Z-stack images and the shortest distance between each spot center and the nearest point of the surface or cell object was calculated based on a 3D distance map. Spots objects (PLA dots) with a distance smaller than 380nm from surfaces (mitochondria) and cell (plasma membrane) objects were considered at a close proximity of these objects. The threshold of 380 nm was used as an estimation of the PLA reaction precision including both primary and secondary antibodies (30nm) plus half the FWHM of the PLA amplification signals (350nm).

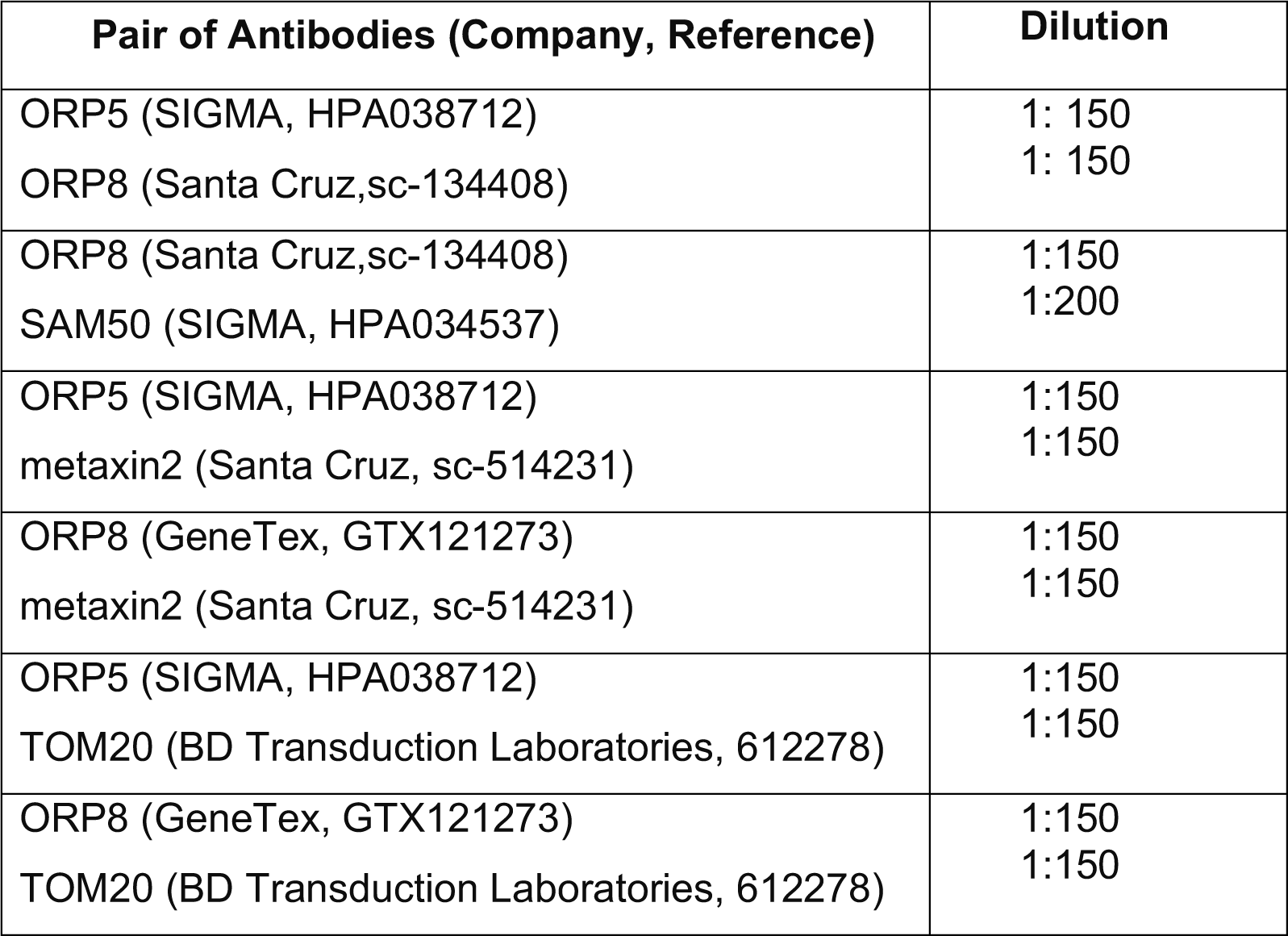

### Electron Microscopy Analysis

#### Conventional EM

For conventional EM, cells grown on 13 mm glass bottom coverslips (Agar Scientific) were fixed with 2.5% glutaraldehyde and 2% PFA in 0.1 M cacodylate, 0.05% CaCl_2_ buffer for 24 hours. After several washes with 0.1 M cacodylate buffer, the cells were postfixed with 1% OsO_4_, 1.5% potassium ferricyanide in 0.1M Cacodylate for 1 hour. After several washes with 0.1 M cacodylate buffer and H_2_O, the cells were stained with 0.5% uranyl acetate for 24 hours. After several washes with H_2_O, the cells were dehydrated in ethanol and embedded in Epon while on the coverslips. Ultrathin sections were prepared, counterstained with uranyl acetate and observed under a MET JEOL 1400 equipped with a Orius High speed (Gatan) camera.

#### HRP Detection

HeLa cells expressing HRP-KDEL were fixed on coverslips with 1.3% glutaraldehyde in 0.1 M cacodylate buffer, washed in 0.1 M ammonium phosphate [pH 7.4] buffer for 1 hour and HRP was visualized with 0.5 mg/ml DAB and 0.005% H_2_O_2_ in 0.1 M Ammonium Phosphate [pH 7.4] buffer. Development of HRP (DAB dark reaction product) took between 5 min to 20 min and was stopped by extensive washes with cold water. Cells were postfixed in 2% OsO_4_+1% K_3_Fe(CN)_6_ in 0.1 M cacodylate buffer at 4°C for 1 hour, washed in cold water and then contrasted in 0.5% uranyl acetate for 2 hours at 4°C, dehydrated in an ethanol series and embedded in epon as for conventional EM. Ultrathin sections were counterstained with 2% uranyl acetate and observed under a FEI Tecnai 12 microscope equipped with a CCD (SiS 1kx1k keenView) camera.

#### Immunogold labelling

HeLa cells were fixed with a mixture of 2%PFA and 0.125% glutaraldehyde in 0.1 M phosphate buffer [pH 7.4] for 2 hours, and processed for ultracryomicrotomy as described previously (Slot & Geuze, 2007). Ultrathin cryosections were single- or double-immunogold-labeled with antibodies and protein A coupled to 10 or 15 nm gold (CMC, UMC Utrecht, The Netherlands), as indicated in the legends to the figures. Immunogold-labeled cryosections were observed under a FEI Tecnai 12 microscope equipped with a CCD (SiS 1kx1k keenView) camera.

| Antibodies (Company, Reference) | Dilution |
| --- | --- |
| GFP (Life technologies, A11122) | 1:100 |
| HA (SIGMA, H3663) | 1:500 |
| PDI (Genetex, GTX30716) | 1:500 |

For the quantification of the number of cristae junction in Epon sections, about 200 mitochondria were analyzed in randomly selected cell profiles and cristae junctions were counted in each of the mitochondria profile and reported as number of cristae/mitochondria profile. All data are presented as mean ±SEM of three experimental replicates.

For the quantification of ER-mitochondria contact sites in HRP-stained Epon sections, the total circumference of each mitochondria and the length of the multiple HRP-positive ER segments closely associated (<30 nm) with them were measured by manual drawing using the iTEM software (Olympus), as in (Galmes et al., 2016, Giordano et al., 2013), on acquired micrographs of HeLa cells for each cell profile, as indicated in the figure legends. Cells were randomly selected for analysis without prior knowledge of transfected plasmid or siRNA. All data are presented as mean (%) ±SEM of three experimental replicates.

For the quantifications of the total mitochondria surface the total circumference of each mitochondria was measured by manual drawing using the iTEM software (Olympus), and data are shown as average of total mitochondrial surface length (μm)/cell ±SD of three experimental replicates.

For the quantification of ORP5 immunogold labeling on ultrathin cryosections, 150 gold particles localized at ER-mitochondria contact sites were counted on acquired micrographs of randomly selected cell profiles at specific ranges of distance from CJ (0-50, 50-100, 100-150, 150-200 nm) in each of three experiments. All data are presented as mean (%) ±SEM of three technical replicates.

### ORP5 and ORP8 ORD domain purification

Escherichia coli BL21DE3 RILP (Invitrogen) cells were transformed with plasmids encoding for GST tagged ORP5 (aa 265-703) or ORP8 (aa 328-767) ORD domains following the manufacturer’s instruction. Bacteria were then grown overnight at 37°C and used to inoculate a large-scale volume (1L). When the OD_600_ reached 0.4, cultures were cooled down and incubated at 18°C until they reached O_D600_ = 0.65. Cells were induced by addition of isopropyl β-D-1-thiogalactopyranoside to a final concentration of 0.1 mM and incubated overnight at 18°C before harvesting. Cells were resuspended in 35 mL binding buffer (1X PBS, 1 mM EDTA, 1 mM DTT, Protease inhibitor) then 250 units of benzonase nuclease (Sigma) were added to the resuspension. Cells were lysed by sonication and the supernatant was recover after 20 min centrifugation at 184 000g and 4°C. Supernatant containing GST tagged proteins was incubated with 2 mL of Glutathione Sepharose 4 fast flow for 1 hour at 4°C under nutation. Beads were washed using a series of wash buffers: 1^st^ (1X PBS, 1 mM EDTA, 1 mM DTT), 2^nd^ HSP-removal buffer (50 mM Tris pH 7.5, 50 mM KCl, 20 mM MgCl_2_, 5 mM ATP) then cleavage buffer (50 mM Tris pH 7.5, 150 mM NaCl, 1 mM EDTA, 1 mM DTT). Cleavage of the GST tag was realized overnight at 4°C using Prescission protease. Untagged proteins were eluted with cleavage buffer, flash frozen and stored at -80°C until lipid transfer assay was performed.

### Liposome preparation

1-palmitoyl-2-oleoyl-sn-glycero-3-phosphocholine (POPC), 1,2-dioleoyl-sn-glycero-3-phosphoethanolamine-N-(cap biotinyl) (Biotinyl Cap PE), 1-palmitoyl-2- (dipyrrometheneboron difluoride)undecanoyl-sn-glycero-3-phosphoethanolamine (TopFluor-PE), 1-palmitoyl-2-(dipyrrometheneboron difluoride)undecanoyl-sn-glycero- 3-phospho-L-serine (TopFluor-PS), 1-palmitoyl-2-(dipyrrometheneboron difluoride)undecanoyl-sn-glycero-3-phosphocholine (TopFluor-PC) were purchased from Avanti Polar Lipids as chloroform solutions.

1 µmol of the appropriate lipid mixtures in chloroform solution was dried in a glass tube for 10 min under a gentle stream of argon, and then for 1 hour under vacuum. The dried lipid films were resuspended in 1 mL of buffer H (25 mM HEPES/KOH, pH 7.7; 150 mM KCl; 10%(v/v) Glycerol) by vigorously vortexing for 30 min at room temperature. Unilamellar liposomes were produced by seven freeze-thaw cycles (30 sec in liquid nitrogen followed by 5 min in a 37°C water bath) and extrusion (at least 21 times) through a polycarbonate filter with 100 nm pore size (polycarbonate membranes from Avanti Polar Lipids). The liposomes were then stored on ice.

### Lipid Transfer assay *in vitro*

The lipid transfer assays were realized with liposomes prepared as described above. The donor liposomes contained 1% mol TopFluor lipids (-PS, -PC or -PE) and 2% mol Biotinyl Cap PE and 97 mol% POPC. The acceptor liposomes contained only POPC. For each reaction, 25 µL of streptavidin-coated magnetic beads (DynabeadsMyOne Streptavidin T1, Invitrogen) were washed in buffer H and mixed with 25 µL of 1 mM donor liposomes. The mixture was incubated for 1 hour at 25°C with intermittent gentle mixing. Bead-bound donor liposomes were then washed, resuspended in 25 µL and mixed with 25 µL of 1 mM acceptor liposomes and 50 µL of buffer H or protein (0.3 µM protein and 2.5 µM TopFluor lipids in the reaction solution). The mixture was incubated at 37°C for 1 hour with intermittent gentle mixing. Supernatant containing acceptor liposomes was recovered after binding of bead-bound donor liposomes to a magnetic rack. TopFluor fluorescence of acceptor and donor liposomes was measured (after solubilization with 0.4% (w/v) n-dodecyl-β-D-maltoside, DDM) in a SpectraMax M5 plate reader (Molecular Device) equilibrated to 30°C (excitation: 450 nm; emission: 510 nm; cutoff: 475 nm; low gain). To confirm that fluorescence was transferred to acceptor liposomes, a fraction of the reaction supernatant – which has not been solubilized with DDM – was floated on a Nycodenz density gradient. 50 µL of supernatant was mixed with 100 µL of buffer H and 150 µL of Nycodenz 80% in buffer H. The solution was transferred to a 0.8 mL Ultra-Clear centrifuge tube (Beckman Coulter) and overlaid with 250 µL of Nycodenz 30% in buffer H and 75 µL of buffer H. The tubes were centrifuged in a SW 55 Ti rotor (Beckman Coulter) at 246,000 g for 4 hours at 4 °C. 50 µL were collected from the top of the gradient and the fluorescence was measured. The percentage of lipids transferred from donor to acceptor liposomes in the presence of ORD5 and ORD8 was determined using the following formula: 100*F_acceptor_/(F_acceptor_+F_donor_).

### Radiometric assay for the conversion of PS to PE *in situ*

#### Radiometric assays without drugs

Hela cells were seeded on 6-well plates and transfected for 48 hours with the non-targeting, ORP5 or ORP8-specific siRNAs specified above by using Oligofectamine (Thermo Fisher Scientific). The cells were then washed and shifted into Hanks balanced salt solution (Gibco) supplemented with a serine-free MEM amino acid mixture and MEM vitamins (Gibco), followed by 18 hours labeling with 2 μl/well L- [^3^H(G)]serine (30.9 Ci/mmol, NET24800, Perkin-Elmer)(Fig. 6c). After the labeling (Fig. 6c) or the chase (Fig. 6d), the cells were scraped into 0.9 ml 2% NaCl per well, a 0.1 ml aliquot was withdrawn for protein analysis with the BCA assay (Thermo Fisher Scientific), and, after adding 50 nmol of unlabeled PS as carrier, the remaining 0.8 ml was subjected to lipid extraction by an acid modification of the Folch method (Kim, Song et al., 2017). After drying, the lipids were resolved in 50 μl CHCl_3_ and applied on Merck TLC Silica gel 60^TM^ plates, followed by separation by using CHCl_3_-methanol-acetic acid-H2O (50:30:8:3.5) as solvent. The PS and PE spots identified from the mobility of standards run on the same plates were scraped into scintillation vials for analysis of [^3^H] radioactivity. The DPM values were corrected for total cell protein, and the ratio of [^3^H] in PE vs. PS calculated.

#### Radiometric assays with drugs

Post 48hr incubation of the cells with transfection complex, media was removed and cell monolayer was washed once with PBS and once with serum-free DMEM. Cells were incubated in a serum-free medium for 12 h. In some experiments, either β-chloro-L-alanine (1 mM) or Hydroxylamine (5 mM) was added to serum-free DMEM, 30 mins prior to pulse, i.e after 11 hr 30 mins serum starvation, cells were starved in serum-free DMEM containing the inhibitor for another 30 mins. Cells were washed twice with PBS, and then incubated for 1 hr at 37 °C, CO_2_ incubator in HBSS media containing 1X MEM amino acids and 1X MEM vitamins containing 7 μCi/ml of [^3^H]serine. After the pulse, the cells were washed twice with PBS and chased for 4 hr in serum-free DMEM at 37 °C. The chase media also contained β-chloro-L-alanine (1 mM) or Hydroxylamine (5 mM). Samples were then washed twice with ice-cold PBS, scraped into 2% NaCl, and the lipids were extracted according to Bligh and Dyer. A fraction of each sample was lysed for protein estimation and blot as described above.

### Mitochondrial respiration assay

Oxygen Consumption rate (OCR) was measured using the XF_p_ Extracellular Flux Analyzer (Seahorse Bioscience Inc.). HeLa cells were seeded on a 6-well plate 3 days before the Seahorse experiment and KD of the proteins of interest was realized 2 days before. The day after KD, HeLa cells transfected with Ctrl, ORP5, or ORP8 siRNAs were plated in a Seahorse XFp 8-mini wells microplate. 20,000 HeLa cells were seeded in each well (except in the blank wells used for the background correction) in 180 μl of culture medium, and incubated overnight at 37 °C in 5% CO2. One day after, the culture medium was replaced with 180 µl of XF DMEM Medium Solution pH 7.4 and then the 8-mini wells microplate was moved in a 37°C non-CO2 incubator before measurement. OCR was determined before drug additions and after addition of Oligomycin (1.5 μM), Carbonyl cyanide 4-(trifluoromethoxy) phenylhydrazone (FCCP, 0.5 μM), and Rotenone/Antimycin A (0.5 μM) (purchased from Agilent). After each assay, all the raw OCR data were analyzed using WAVE software.

### Quantifications and Statistical Analysis

Statistical analysis was performed with Microsoft Excel or GraphPad Prism 9.0. The data were presented as mean ± SEM. The n, indicated in the figures and figure legends, represent the total number of cells analyzed in three or more biological replicates, as stated in the figures legend. Statistical significance of two data sets were determined by unpaired student’s *t*-test, with * p<0.05, ** p<0.01, ***p<0.001 and **** p<0.0001.

## ACKNOWLEDGEMENTS

We thank Dr V. Kozjak-Pavlovic and P. Somerharju for kindly sharing reagents and advices with us. We also thank Dr. R. Legouis, Dr. Emmanuel Culetto for discussion and for critically reading the manuscript; Claire Boulogne, Sandrine Lécart and Remi Le Borgne for help and consultation with microscopy experiments; Valentin Guyard, Blandine Bourigault, Riikka Kosonen and Liisa Arala for technical assistance. The present work has benefited from Imagerie-Gif core facility supported by I’Agence Nationale de la Recherche (ANR-11-EQPX-0029/Morphoscope, ANR-10-INBS-04/FranceBioImaging; ANR-11-IDEX-0003-02/ Saclay Plant Sciences). For the Immuno-EM and HRP-KDEL EM analysis/quantifications we acknowledge the ImagoSeine facility, member of the France BioImaging infrastructure supported by grant ANR-10-INBS-04 from the French National Research Agency. We also thank Jean-Michel Camadro and Thibault Leger (IJM proteomics platform) for proteomics analysis. This work was supported by the ANR Jeune Chercheur (ANR0015TD), the ATIP-Avenir Program, the FSER (FRM n°206548) and the FVA (eOTP:669122 LS 212527) to F.G.; the Academy of Finland (grants 285223 and 322647), the Sigrid Juselius Foundation, the Finnish Foundation for Cardiovascular Research and the Magnus Ehrnrooth Foundation to V.M.O.; the “Association Française contre les Myopathies” (AFM Research grant 20123) and the ITMO-Inserm Plan Cancer 2014- 2019 to D.T.

## AUTHOR CONTRIBUTIONS

FG conceived and supervised the work. VO designed and supervised the radiometric assays for PS-to-PE conversion and the expression analysis by RT-PCR. DT designed and supervised the *in vitro* lipid transfer assays. LR performed and analyzed the cell experiments including cell imaging and Seahorse analysis. VC performed and analyzed most of the cell experiments including immunofluorescence for endogenous proteins and Duolink. LR, AH, and FG performed and analyzed the EM experiments. CS participated to the setting up of the Seahorse experiments. CS, AH and DT performed and analyzed the *in vitro* lipid transfer assays. AA, EJ and AK performed and analyzed the radiometric assays for PS-to-PE conversion. EM, JD and JS performed MS-lipidomic analysis. RLB provided tools and techniques for Duolink imaging analysis. JN and NEK provided technical help in western blot analysis and generated some of the constructs for mammalian cell expression. VC, LR and FG wrote the manuscript and all authors commented on the manuscript.

## DATA AVAILABILITY SECTION

This study includes no data deposited in external repositories.

---

**Figure S1.**
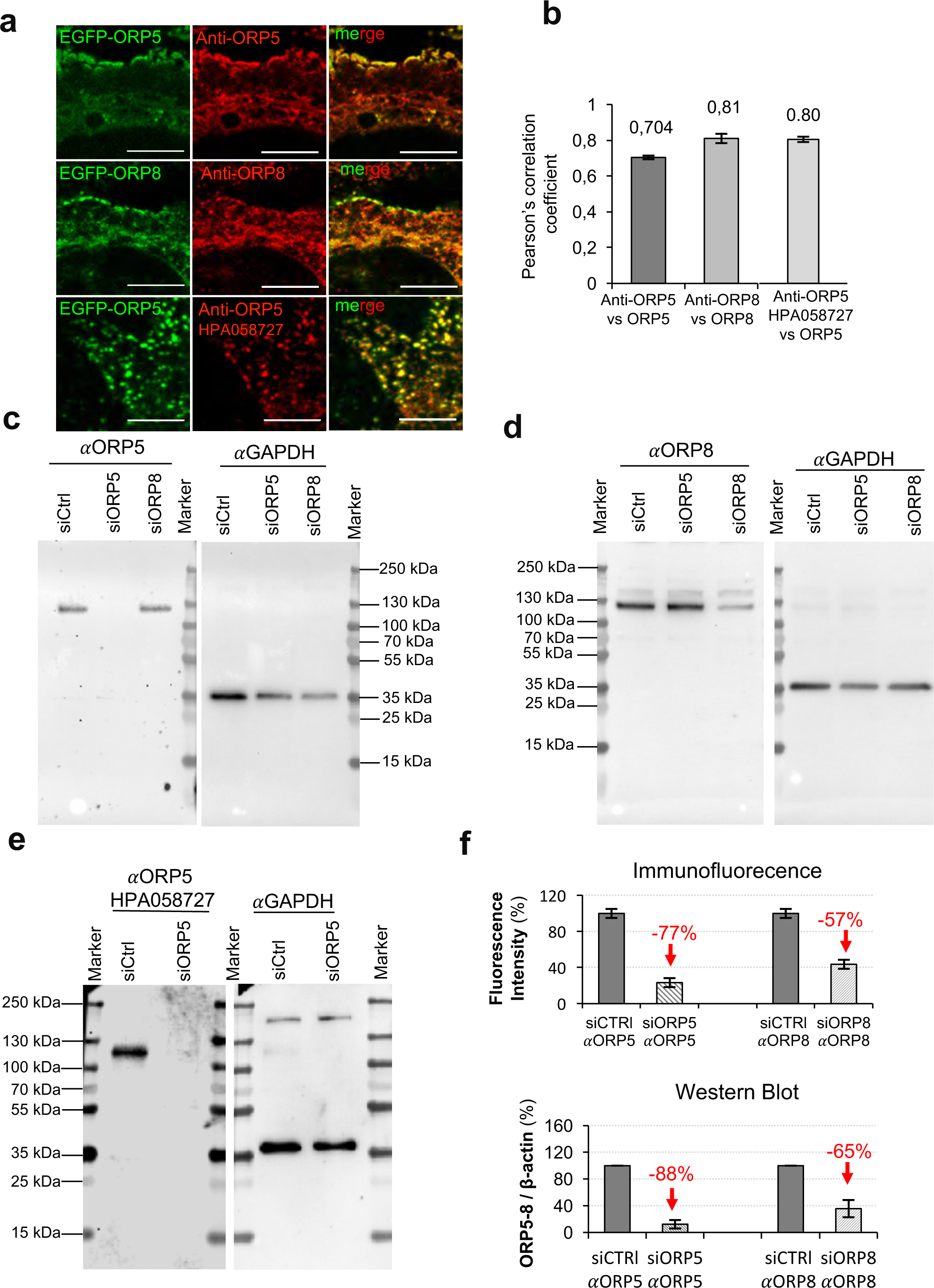
Antibodies anti-ORP5 and anti ORP8 specifically bind ORP5 and ORP8, respectively. (a) Confocal images of a region of HeLa cell transfected with EGFP- ORP5 (green) or EGFP-ORP8 (green) and immunostained using anti-ORP5 (red) or anti- ORP8 (red) antibodies. Scale bar, 5 μm. (b) Quantifications of the colocalization (Pearson’s factor) of EGFP-ORP5 or EGFP-ORP8 with anti-ORP5 or anti-ORP8. Bars indicate mean values ±SEM. Cells analyzed for sample: n=5. (c-e) Western blot analysis using primary antibodies against ORP5 and ORP8 in protein samples obtained from HeLa cells treated with scrambled siRNA (siCtrl) and from Hela cells treated with siRNA targeting ORP5 (siORP5) or ORP8 (siORP8). Images show the full membranes which were firstly incubated with anti-ORP5 or anti-ORP8, and then with anti-GAPDH. (e) Analysis of ORP5 and ORP8 expression in siCtrl and siORP5 and siORP8, respectively, by immunofluorescence and Western blot.

**Figure S2.**
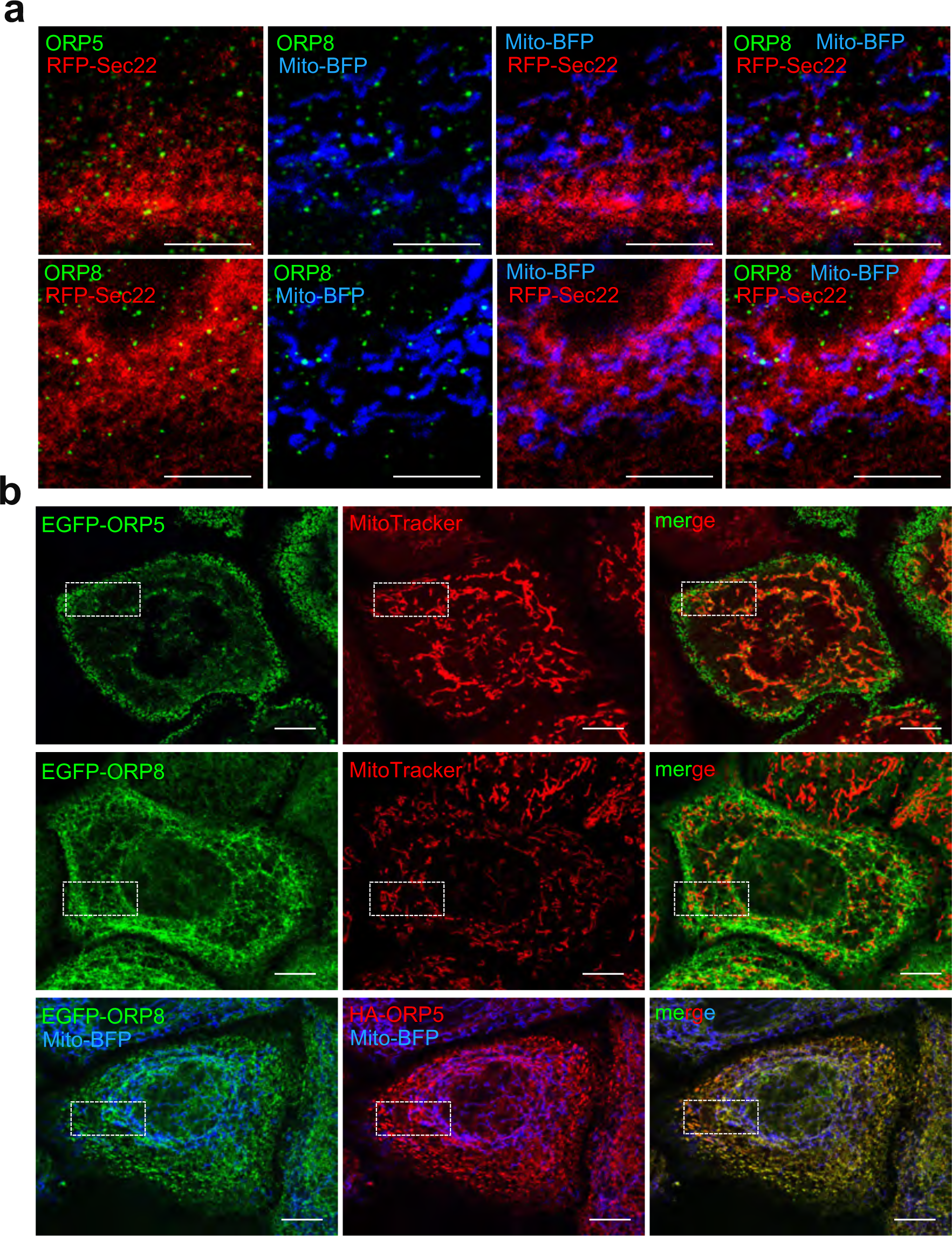
Endogenous and co-overexpressed ORP5 and ORP8 co-localize at ER-mitochondria contacts. (a) Confocal micrographs of regions of HeLa cell transfected with RFP-Sec22b (red) together with Mito-BFP (blue) and immunostained for anti-ORP5 (green) or anti-ORP8 (green) antibodies. Scale bar, 5 μm. (b) Confocal micrograph of a HeLa cell transfected with EGFP-ORP5 (green), EGFP-ORP8 (green) or EGFP-ORP8 (green) + HA-ORP5 (anti-HA, red), together with Mito-BFP (blue). The boxed areas are magnified in Fig 1g. Scale bar, 10 μm.

**Figure S3.**
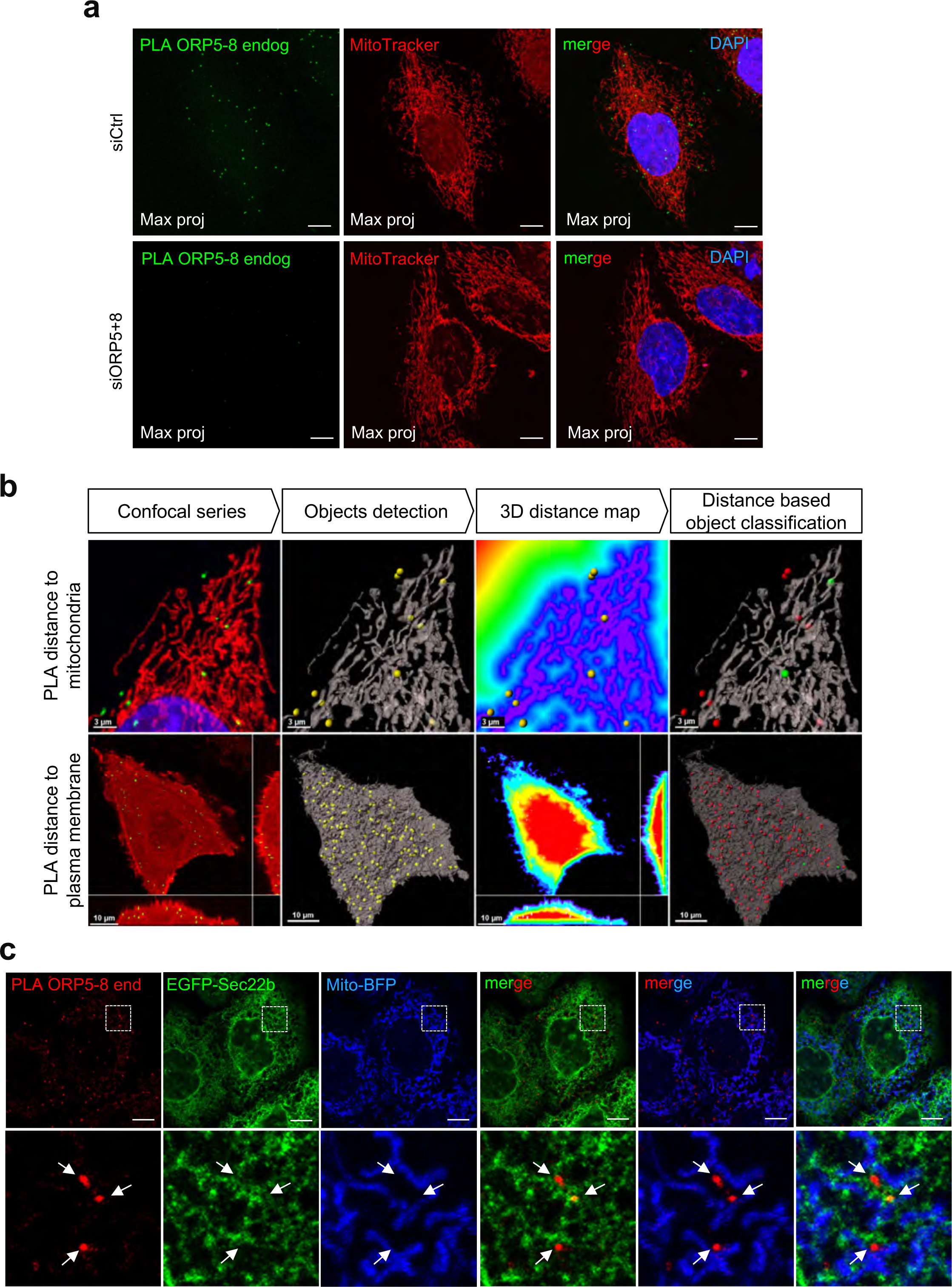
Endogenous ORP5-ORP8 interaction at ER-mitochondria contact sites. (a) Confocal images showing endogenous interaction of ORP5-8 by Duolink PLA (green) in Ctrl (siCtrl), and ORP5+ORP8 (siORP5+8) knockdown HeLa cells labeled with MitoTracker (red) to stain mitochondria. Images are presented as maximum projection of all layers. Insets show magnifications of the boxed regions. Scale bar, 10 μm. (b) Workflow for the identification of PLA signals in close proximity to the mitochondria or plasma membrane. First the confocal stacks are segmented to identify the PLA foci (spots), the mitochondria network (surfaces), and the plasma membrane (cells). Then 3D distance maps are computed towards the outside of the surfaces (mitochondria) or inside the cells allowing the measurement of the distance of each spot from the closest mitochondria or membrane. Finally, PLA spots are classified into two population (red and green) based on a proximity threshold of 380nm established on the precision of the detection system. (c) Confocal micrographs of HeLa cell transfected with GFP-Sec22b (green) and Mito-BFP (blue) and showing endogenous interaction of ORP5-8 by Duolink PLA (red). Images are presented as superposition of two layers. Scale bar, 5 μm.

**Figure S4.**
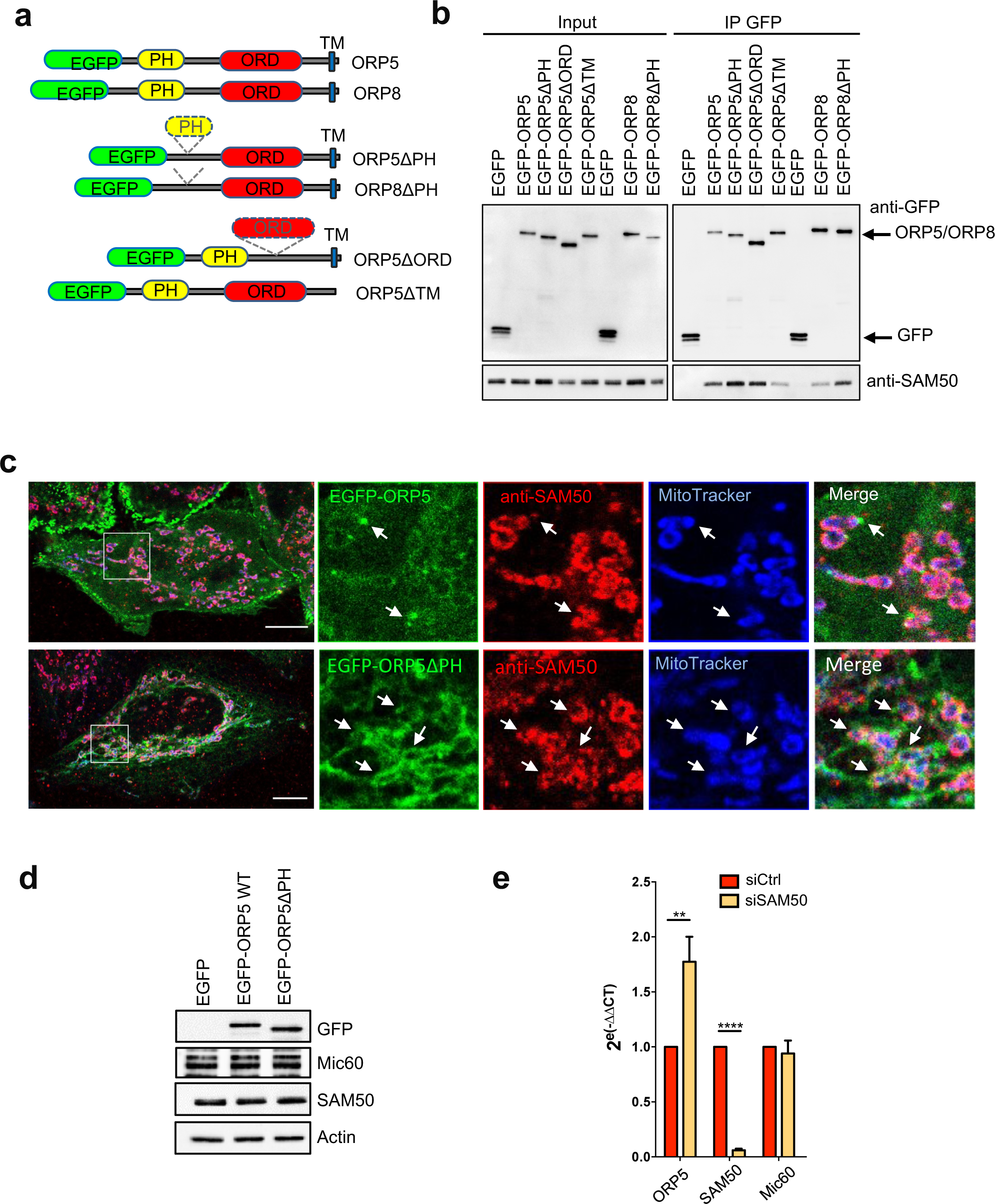
Localization of ORP5 near SAM50-labeled mitochondria and effect of SAM50 KD on ORP5 and Mic60 transcription. (a) ORP5 and ORP8 full-length and mutant constructs used in Fig. 4a-b and in in Fig. S4b-c. (b) EGFP-ORP5, EGFP- ORP5ΔPH, EGFP-ORP5ΔORD, EGFP-ORP5ΔTM, EGFP-ORP8, EGFP-ORP8ΔPH or EGFP alone were transfected in HeLa cells then immuno-precipitated from lysates and analyzed by western blot using antibodies against GFP (for ORP5 or ORP8) or against SAM50. (c) Confocal micrograph of a HeLa cells transfected with EGFP-ORP5 or EGFP- ORP5ΔPH (green) and Mito-BFP (blue) and stained with anti-SAM50 (red) antibody. Insets show magnifications of the boxed regions. Scale bar, 10 μm. (d) WB analysis showing GFP (EGFP-tagged constructs), Mic60, SAM50 and Actin levels in protein lysates from HeLa cells transfected with either EGFP (Control) or with EGFP-ORP5 or EGFP-ORP5ΔPH constructs. (e) Quantitative RT-PCR analysis of ORP5, SAM50 and Mic60 in) values of tested genes and those of reference gene (SDHA) in SAM50 knockdown cells versus control HeLa cells. SAM50 knockdown does not alter Mic60 RNA levels and it induces an increase in ORP5 transcription. y axis: 2^(-ΔΔCt)^ value represents differences between the mean Ct (Cycle threshold). Statistical analysis: unpaired student’s t-test , **P<0.01, ****P<0.0001.

**Figure S5.**
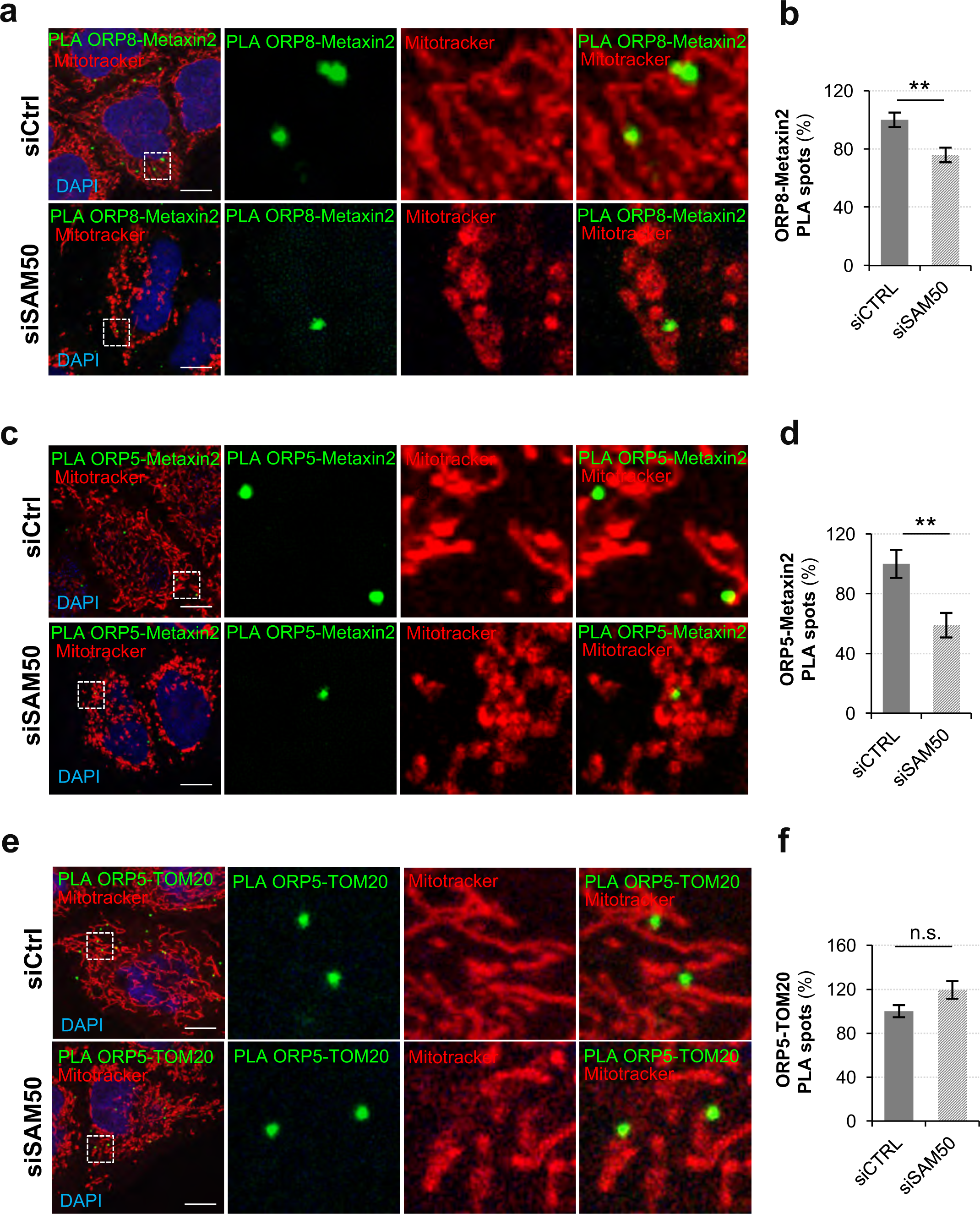
SAM50 regulate PLA interactions of ORP5-metaxin2 and ORP8- Metaxin2, but not ORP5-TOM20. (a-c) Representative confocal images showing endogenous ORP5-metaxin2, ORP5-TOM20 and ORP8-metaxin2 PLA interactions (green), mitochondrial network (MitoTracker, red) and nuclei (DAPI, blue) in control (siCtrl) and SAM50 (siSAM50) knockdown HeLa cells. Images are displayed as single layers. Scale bar, 10 μm. (d-f) Quantitative analysis of endogenous endogenous ORP5-TOM20 (n= 49 cells siCtrl, and n= 49 cells siSAM50) ORP5-TOM20 (n= 51ccells siCtrl, and n=52 cells siSAM50) and ORP5-metaxin2 (n= 35 cells siCtrl, and n= 24 cells siSAM50) PLA interactions in control and SAM50 (siSAM50) knockdown HeLa cells. Data are shown as mean values ±SEM. Statistical analysis were performed using unpaired student’s t-test, with **, P<0.01, n.s., not significant.

**Figure S6.**
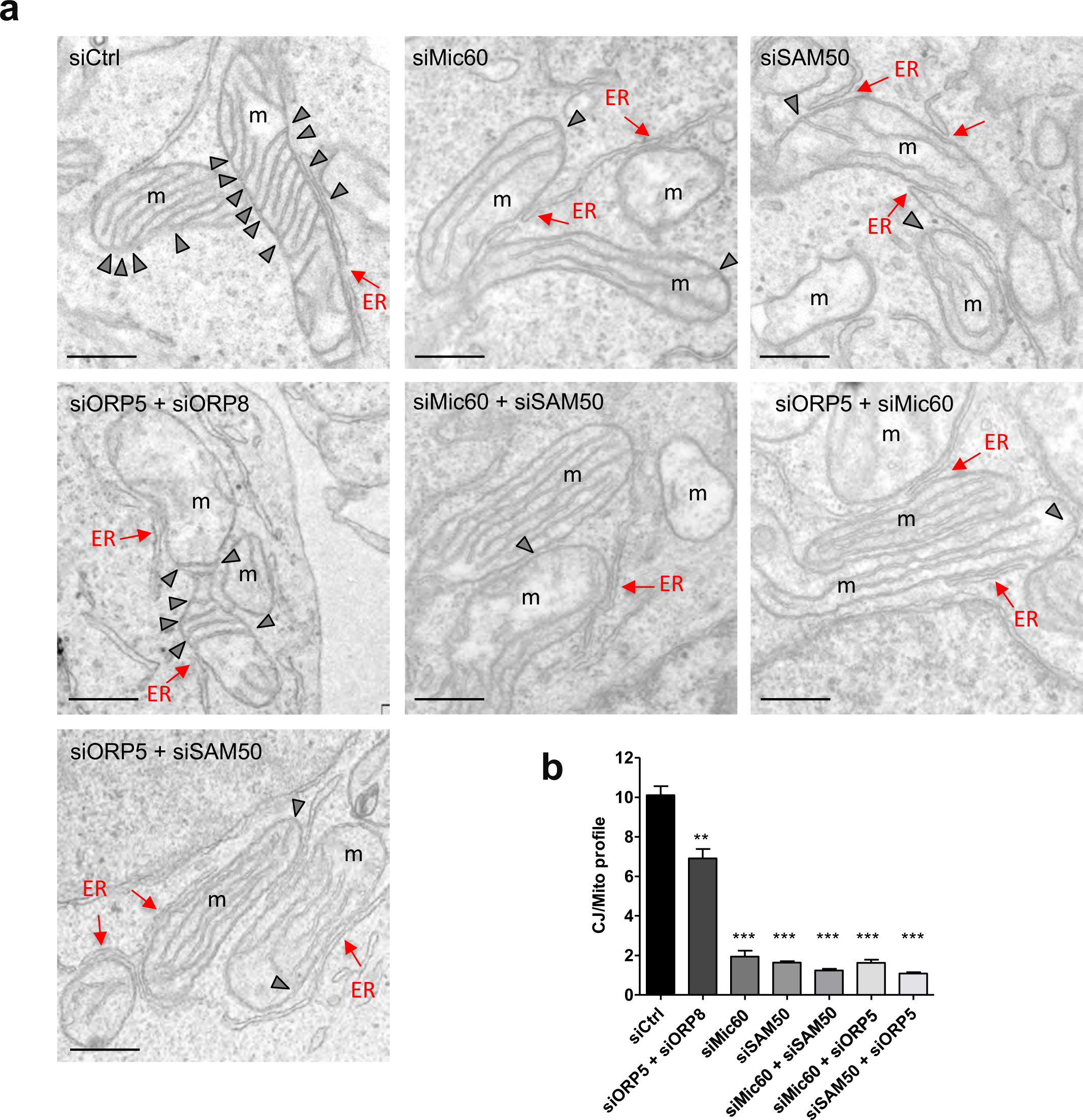
Cristae Junctions are altered upon SAM50, Mic60 and ORP5+8 knockdowns. (a) Representative EM micrographs showing the morphology of mitochondria in HeLa cells treated with siRNA against ORP5, ORP8, Mic60 and SAM50. Scale bar, 200 nm. Red arrows indicate ER elements in contact with mitochondria; arrowheads indicate CJ; m, mitochondria. (b) Quantifications of the number of CJ per mitochondria profile in the indicated siRNA conditions. % of CJ ±SEM, n = 170-260 mitochondria. Statistical analysis: unpaired student’s t-test , **P<0.01, ***P<0.001.

**Figure S7.**
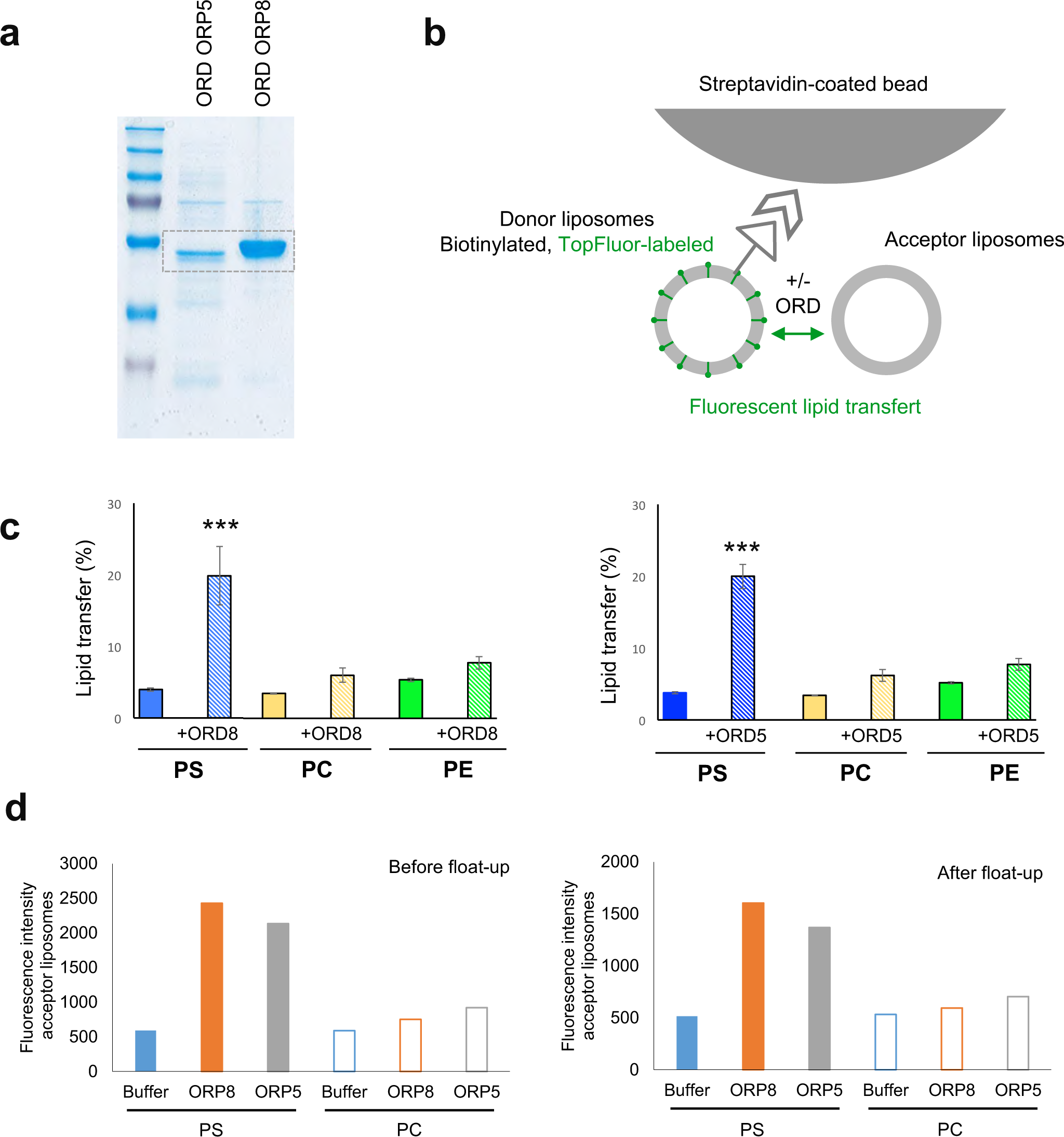
ORP5 and ORP8 ORD domains mediate PS transfer between liposomes in vitro. (a) Coomassie stained SDS-PAGE of the recombinant ORD domain of ORP5/8 proteins purified from BL21DE3 RILP cells. (b) Schematic cartoon of the in vitro assay used to study ORP5/8 ORD-mediated lipid transport between liposomes. (c) Donor liposomes containing fluorescent lipids (97 mol% POPC, 1 mol% TopFluor-PS, - PC or -PE and 2 mol% of biotinylated-PE) and pre-bound to streptavidin beads were mixed at a 1:1 molar ratio with acceptor liposomes (100 mol% POPC) in the presence or absence of ORP5 or ORP8 ORD domains (250 µM of donor and acceptor liposomes and 0,3 μM of proteins in the reaction solution). The increase in fluorescence in the acceptor liposomes, which remain unbound in the solution, was measured after 1 hour of incubation at 37°C. Data are presented as % of transferred lipid ±SEM and are the mean of six independent experiments. Statistical analysis: unpaired student’s t-test , ***P<0.001. (d) Results of a lipid transfer experiment performed as in (c) and presented as the fluorescence intensity of acceptor liposomes before (left panel) or after (right panel) their floatation on a Nycodenz gradient to confirm that fluorescence comes from the liposomes membrane.

**Figure S8.**
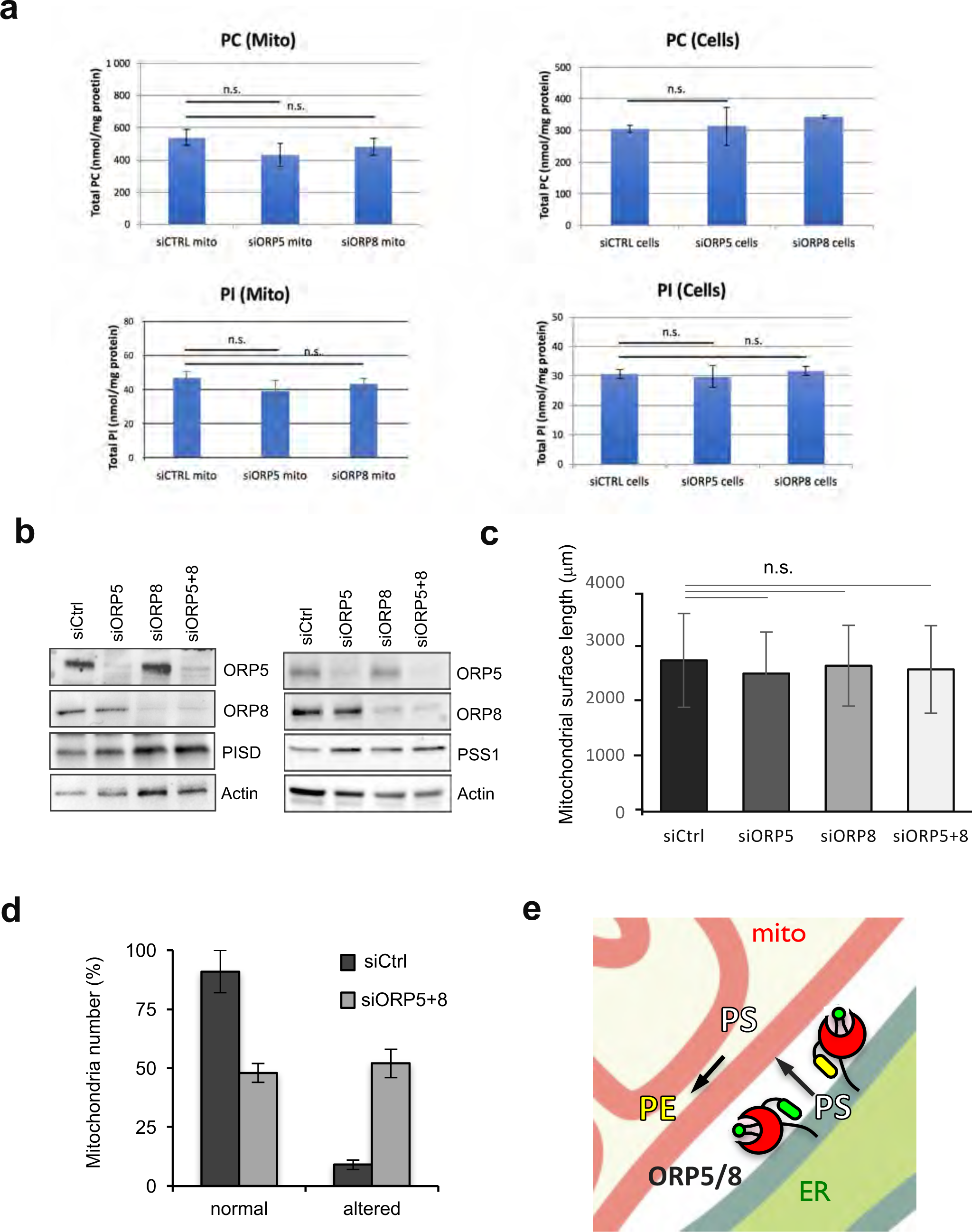
ORP5/8 specifically regulate levels of PS-derived mitochondrial PE and mitochondria morphology but not PSD1 and PSS1 protein levels in situ. (a) MS- based quantification of PC and PI content (nmol/mg protein) of mitochondria isolated from Ctrl, ORP5 or ORP8 knockdown cells and of Ctrl, ORP5 or ORP8 knockdown intact cells. Data are shown as mean of three independent replicates ±SEM. Statistical analysis: unpaired student’s t-test. n.s. not significant. (b) WB analysis showing ORP5, ORP8, PSD1, PSS1 and Actin levels in protein lysates from HeLa cells treated with either Ctrl siRNAs or with siRNAs against ORP5 or/and ORP8. (c) Quantifications of the total mitochondria surface length. Data are shown as mean of total mitochondrial surface length (mm)/cell ±SD, n = 20 cell profiles and ±900 mitochondria. Statistical analysis: unpaired student’s t-test, n.s. not significant. (d) Quantifications of the number of mitochondria with aberrant cristae morphology in the indicated siRNA conditions. Data are shown as % of mitochondria ±SEM, n = 20 cell profiles and ±700 mitochondria. Statistical analysis: unpaired student’s t-test, **P<0.01 compared to siCtrl. (e) Schematic representation of non-vesicular PS transfer mediated by ORP5/8 at ER-mitochondria contact sites. PS is transported to mitochondrial membranes where it is rapidly converted into mitochondrial PE.

**Figure S9.**
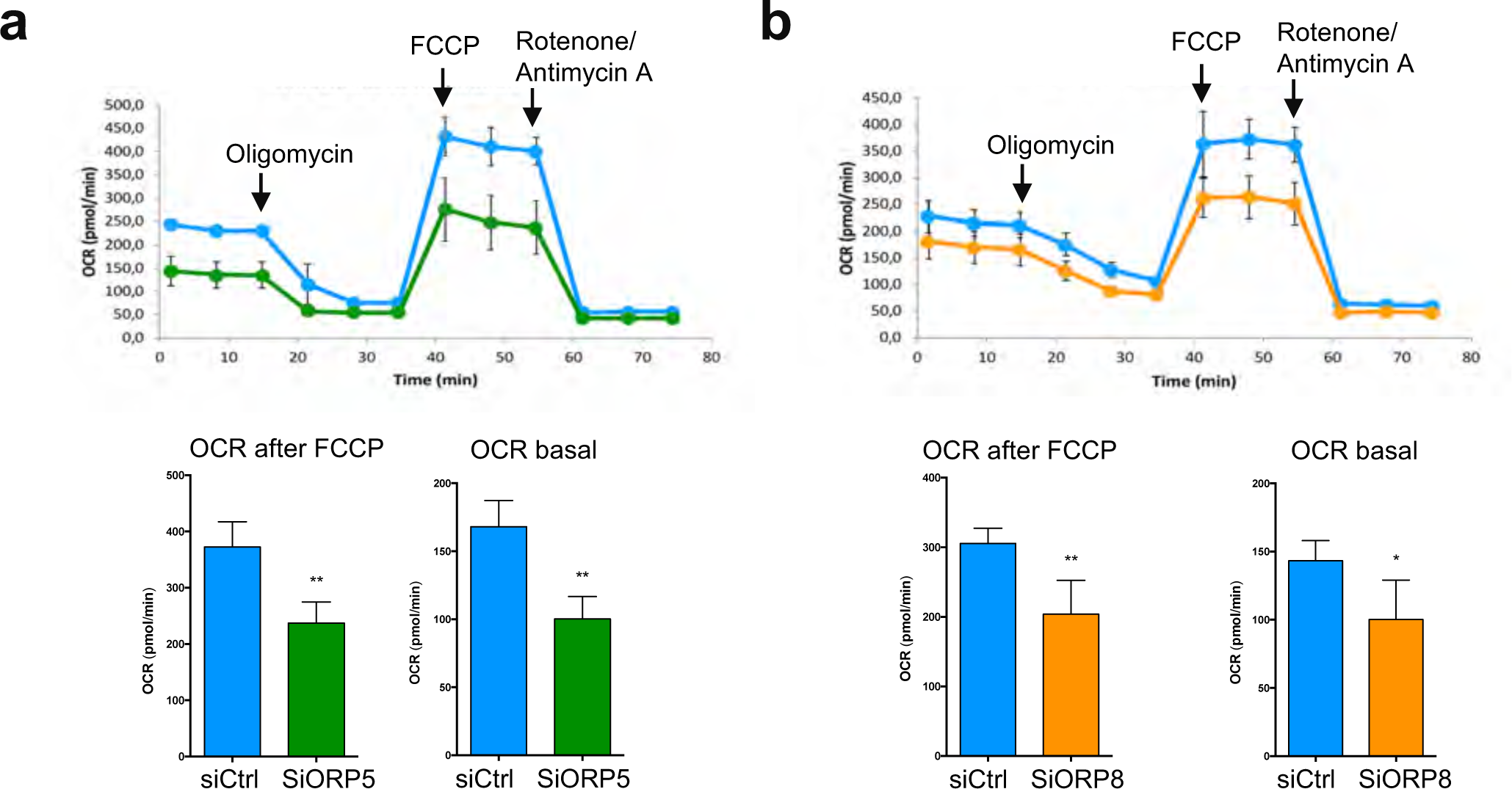
ORP5 and ORP8 knockdowns affect mitochondria respiratory function. (a-b) Mitochondrial oxygen consumption rate (OCR) measured in Ctrl and ORP5 (a) or ORP8 (b), siRNA-treated HeLa cells. OCR trace was obtained by sequential measurement of basal OCR (OCR_BAS_), OCR after the addition of Oligomycin, OCR after the addition of FCCP (OCR_FCCP_) and OCR after the addition of Rotenone/Antimycin A. Note the reduced OCR in siORP5 and siORP8 cells compared to Ctrl siRNA cells. Error bars denote ±SEM. Data shown in the bar charts are the mean of 4 independent repeats (n=4). Statistical analysis: unpaired student’s t-test, *P<0.05, **P<0.01 compared to Ctrl.

